# Computational design of nanoscale rotational mechanics in *de novo* protein assemblies

**DOI:** 10.1101/2021.11.11.468255

**Authors:** A. Courbet, J. Hansen, Y. Hsia, N. Bethel, YJ. Park, C. Xu, A. Moyer, S.E. Boyken, G. Ueda, U. Nattermann, D. Nagarajan, D. Silva, W. Sheffler, J. Quispe, N. King, P. Bradley, D. Veesler, J. Kollman, D. Baker

**Affiliations:** Department of Biochemistry, University of Washington, Seattle, USA; Institute for Protein Design, University of Washington, Seattle, USA; Howard Hughes Medical Institute, University of Washington, Seattle, WA 98195; Division of Public Health Sciences, Fred Hutchinson Cancer Research Center, Seattle, USA

## Abstract

Natural nanomachines like the F_1_/F_0_-ATPase contain protein components that undergo rotation relative to each other. Designing such mechanically constrained nanoscale protein architectures with internal degrees of freedom is an outstanding challenge for computational protein design. Here we explore the *de novo* construction of protein rotary machinery from designed axle and ring components. Using cryoelectron microscopy, we find that axle-ring systems assemble as designed and populate diverse rotational states depending on symmetry match or mismatch and the designed interface energy landscape. These mechanical systems with internal rotational degrees of freedom are a step towards the systematic design of genetically encodable nanomachines.

**One-Sentence Summary:** Computationally designed self-assembling protein rotary machines sample internal degrees of freedom sculpted within the energy landscape.

---

Intricate protein nanomachines in nature have evolved to process energy and information by coupling biochemical free energy to mechanical work. Among the best studied and most sophisticated is the F_1_/F_0_-ATPase which consist of an axle component surrounded by a ring-like component that undergo constrained dynamic rotary motion relative to each other, mediating the synthesis of ATP using the energy stored in a proton gradient(*1*). Inspired in part by Feynman’s 1959 lecture suggesting nanotechnology as the means to leverage the properties of materials at the molecular scale(*2*), there has been a growing interest in the creation of synthetic nanomachines(*3,4*). Synthetic chemists were the first to design technomimetic molecules with mechanically coupled parts(*5–7*). Nucleic acid nanotechnologies have more recently been used to construct rotary systems(*8*). The design of dynamic protein mechanical systems is of great interest given their richer functionality, but while recent advances in protein design now enable the generation of increasingly sophisticated static nanostructures and assemblies(*9–17*), the complex folding and diversity of non-covalent interactions has thus far made this very challenging(*18*).

We set out to explore the design of protein mechanical systems through a first-principle, bottom-up approach that decouples operational principles from the complex evolutionary trajectory of natural nanomachines. Sampling of the folding landscape for both structural and dynamic features is computationally expensive, and hence we decided on a hierarchical design approach with steps that can be tackled in turn: (*i*) the *de novo* design of stable protein building blocks optimized for assembly into constrained mechanical systems, (*ii*) the directed self-assembly of these components into hetero-oligomeric complexes, (*iii*) the shaping of the multistate energetic landscape along mechanical degrees of freedom (DOF) and (*iv*) the coupling of chemical or light energy to rotation or other motion. In this paper, as a proof of concept we aim to assemble a *simple machine* or kinematic pair(*19,20*) at the nanoscale, and focus on steps i-iii to design mechanically constrained heterooligomeric protein systems that undergo brownian rotary motion. We start from a rotary machine blueprint (**fig. 1A**) in which, similar to natural rotary systems, the features of the rotational energy landscape are determined by the symmetry of the interacting components, their shape complementarity and specific interactions across the interface.

**Fig. 1:**
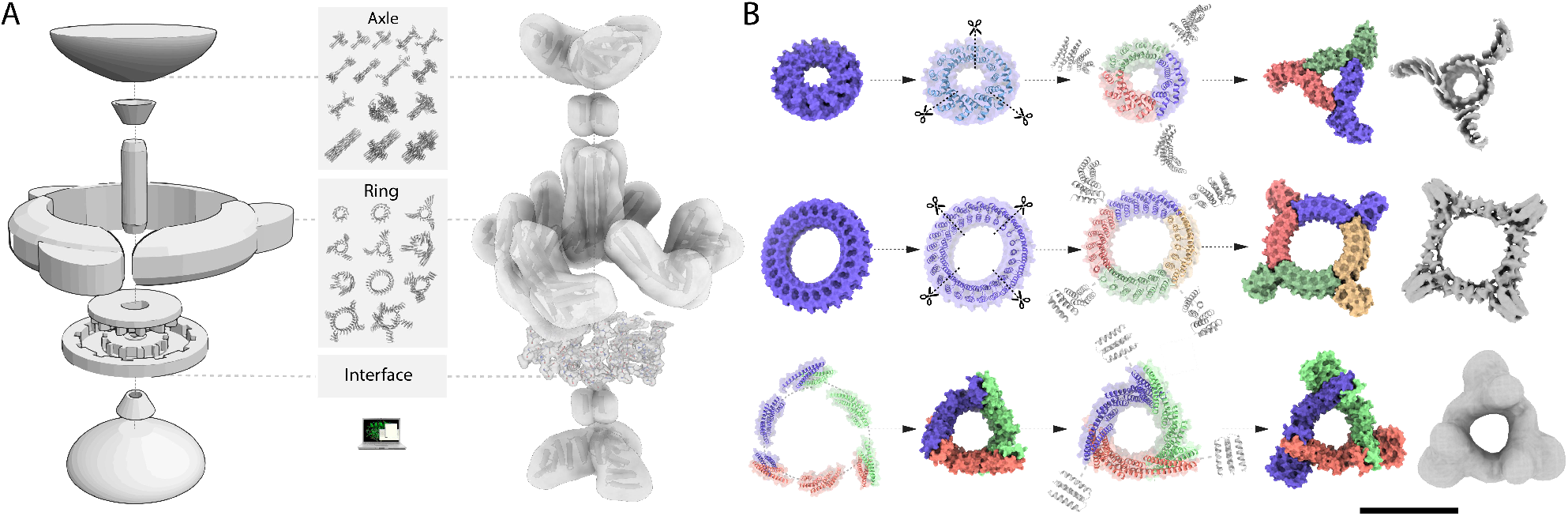
Overview of rotary machine assembly and ring design approaches. (**A**) (Left) A blueprint of a simple rotary machine consisting of an assembly of an axle and a ring, mechanically constrained by the interface between the two; (Middle) Systematic generation of a structurally diverse library of machine components through computational design. The design of the interface between axle and ring mechanically couples the components by providing control on the rotational energy landscape and directing assembly; (Right) Example of hierarchical design and assembly of a protein rotary machine from axle and ring components, here a D3 axle and C3 ring, and interacting interface residues. Cyclic DHRs or *wheels* are fused to the end of the axle and ring components to increase mass, provide a modular handle and a rotation dependent structural signature. (**B**) Hierarchical design strategies for ring components (Top) A single chain C1 symmetric and internally C12 symmetric alpha-helical tandem repeat protein is split into three subunits, and each is fused to DHRs via helical fusion (HelixFuse) to generate a C3 ring with an internal diameter of 28Å. The 6.5Å cryoEM electron density (shown in grey) shows agreement with the design model (monomer subunits colored by chain); (Middle) A single chain C1 symmetric and internally C24 symmetric alpha-helical tandem repeat protein is split into 4 subunits and each is fused to DHRs to generate a C4 ring with an internal diameter of 57Å. The 5.9Å cryoEM electron density (shown in grey) shows agreement with the design model (monomer subunits colored by chain); (Bottom) Heterooligomeric helical bundles and DHRs are fused using HelixDock and HelixFuse and assembled into a higher-ordered closed C3 structure through helical fusion (WORMS), after which another round of helical fusion protocol (HelixFuse) is used to fuse DHRs to each subunit, to generate a C3 ring with an internal diameter of 41Å. The negative stain electron density (shown in grey) shows agreement with the design model (monomer subunits colored by chain). Scale bar: 10nm

## Computational design of protein rotary machine components

We set out to design *de novo* a library of stable protein components with shapes, fold and symmetry specifications suitable for integration into rotationally constrained assemblies. We first sought to design ring-like protein topologies with a range of inner diameter sizes capable of accommodating an axle-like binding partner in the center (**fig. 1B**). In a first design approach, we started from *de novo* designed alpha-helical tandem repeat proteins(*21*), which were redesigned to be C1 single chain structures or symmetric C3 or C4 homooligomers. In a second approach, we used a hierarchical design procedure based on architecture-guided rigid helical fusion(*12*) to build C3 and C5 cyclic symmetric ring like structures by modularly assembling via rigid fusion *de novo* helical repeat proteins (DHRs) and helical bundle heterodimers. To facilitate experimental characterization by optical and electron microscopy, we increased the radius and total mass of the designs by fusing another set of DHRs at the outer side of the rings, generating arm-like extensions (**fig. 1A-B**). Synthetic genes encoding these designs (12xC3s, 12xC4s, 2xC5s) were synthesized and the proteins expressed in *E. coli*. All designed proteins were soluble after purification by nickel-nitrilotriacetic acid (Ni-NTA) and ~23% (6/26) had appropriate monodisperse size exclusion chromatography (SEC) profiles that matched the expected theoretical elution profile for the oligomerization state. These designs were further examined using small-angle X-ray scattering (SAXS)(*22,23*), negative stain electron microscopy or cryoelectron microscopy (cryoEM) (**fig. S1, fig S2**). For the R82 C3 ring, SAXS data was consistent with the computational model and we were able to determine using cryoEM a 6.5Å 3D reconstruction which was very close to the design model (**fig. 1B, fig. S2-4, Table S1**). Additional designs of the same topology (R14 and R76) were characterized by SAXS and showed similar profiles, and hence likely have the same oligomeric state and overall structure (**fig. S2**). A C3 ring with larger inner diameter and different topology (R113) was characterized using negative stained EM, yielding a low resolution 3D reconstruction consistent with the design model (**fig. 1B, fig. S2**). For a C4 design highly expressed in *E. coli*, we obtained a ~5.9Å cryo electron density map revealing a structure nearly identical to the design model (**fig. 1B, fig. S2, fig. S5, Table S1**). Negative stain EM of a C5 ring yielded a low resolution 3D map consistent with the design model (**fig. S2**).

We next sought to design high aspect ratio protein folds, or *axles*, onto which the ring-like designed protein could be threaded. In a first approach, single helix protein backbones were parametrically generated, and then D2, D3 or D4 dihedral symmetry was imposed to produce self-assembling dihedral homooligomers consisting of interdigitated single helices (**fig. 2A**). Two helices were placed roughly colinearly along the z axis but at different distances from it, their superhelical parameters were sampled using the Crick-generating equations(*24*), and those for which imposition of dihedral symmetry generated closely packed structures were connected with a linking helix (see “Computational design methods” in the supplementary materials). Rosetta HBNet(*25*) was then used to install hydrogen bond networks with buried polar residues between the helices (4, 6, or 8 for a D2, D3 and D4 respectively) to generate homooligomeric interfaces with the high level of specificity needed for dihedral assembly. The sequence of the rest of the homooligomer (surface residues and the hydrophobic contacts surrounding the networks) was then optimized while keeping the networks constrained during RosettaDesign as described previously(*25*). Last, in order to increase the total mass, diversify the shape as well as increase the modularity of axles, each helix of the best-scoring designed dihedral homooligomers was connected at either the C or N terminus to an outer helix belonging to *de novo* cyclic homooligomer *wheels of* matching symmetry (*i.e.* Cn -> Dn), through a short helical fragment sampled and designed using Rosetta Remodel, to finally produce full axle homooligomers. In a second approach, *de novo* cyclic homooligomers were selected(*15*) and Rosetta BlueprintBuilder(*26*) was used to generate interdigitated helical fragments of varying length and topology which were computationally extensively sampled at the N or C terminus in order to direct the assembly into dihedral homooligomers (**fig. 2B**, see “Computational design methods” in the supplementary materials). In a third approach, cyclic homotrimer backbones consisting of helical hairpin monomer topologies with inner and outer helices that were previously parametrically generated(*25*) were circularly permuted by re-looping terminis using the Rosetta ConnectChainsMover, placing terminis in the middle of the outer helices, and elongating inner helix heptad repeats to generate C3 homooligomers which 3 inner helix form an accessible surface for further rotary machine design (**fig. 2C)**.

**Fig. 2:**
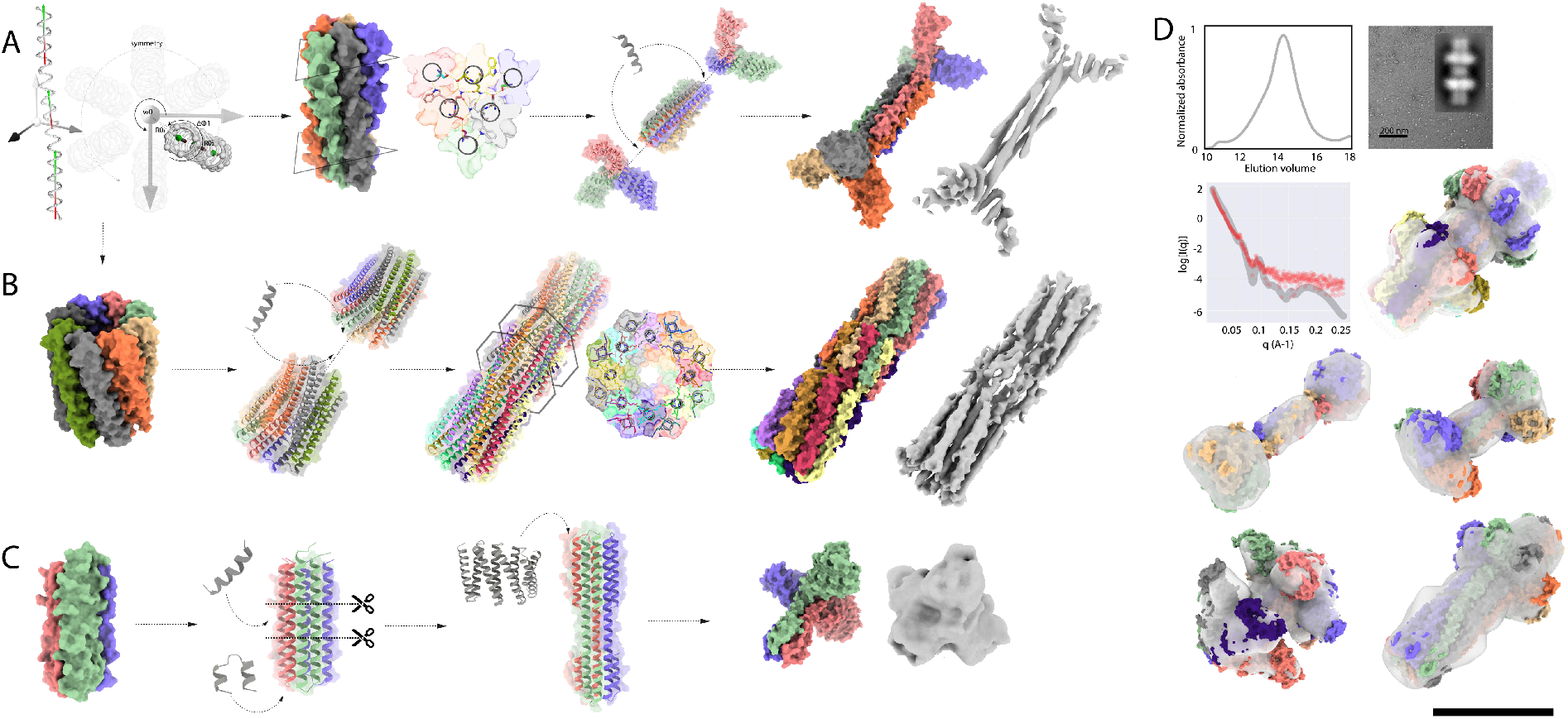
Design of axle machine components. (**A**) Hierarchical design of a D3 symmetric homohexamer axle (1552_1na0C3_int2_11). Parametric design of interdigitated helices in D3 symmetry is achieved by sampling supercoil radius (R_1_,R_2_), helical phase (Δφ_1-1_, Δφ_1-2_), supercoil phase (Δφ_0-1_,Δφ_0-2_) of two helical fragments, and the *z*-offset (Z_off_), and supercoil twist (ω_0_). The interface is designed using the HBNet protocol to identify hydrogen-bond networks spanning the 6 helices mediating high-order specificity. The design is then fused to C3 homotrimers using RosettaRemodel. The 4.2Å cryoEM electron density is consistent with the design model (**B**) Hierarchical design of a D8 axle (D8A_1615). Starting from a parametrically designed C8 homohexamer, interdigitated helical extensions are sampled using Rosetta BluePrintBuilder and hydrogen bond networks identified using HBnet while sampling rotation and translation in D8 symmetry using Rosetta SymDofMover. The 5.9Å cryoEM electron density shows close agreement with the design model; (**C**) Hierarchical design of a C3 homotrimer axle (A15.5). A parametrically designed C3 homotrimer is circularly permutated and an extra heptad repeat is added to increase the aspect ratio, after DHRs are fused to each subunit using Hfuse. The negative stain electron density is consistent with the design model (**D**) Additional axle designs (Top) Representative SEC, SAXS and negative stain EM profile corresponding to a D8 design (D8_6_49). The SAXS trace (red) is similar to the computed trace from the model (grey); (Bottom) Design models for D2_1119_7_tj81C2_V39_6, DC4G1_178, D5_57C, and C8D8_6_49 overlaid with experimental 3D electron density. Model monomer subunits are colored by chain, and electron densities are shown as grey surfaces. Scale bar: 10 nm

Synthetic genes encoding axle designs generated from the three approaches (12xC3s, 12xC5s, 12xC8s, 6xD2s, 12xD3s, 6xD4s, 6xD5s, 12xD8s) were obtained and the proteins were expressed in *E. coli*. The designed proteins that were well-expressed, soluble, and readily purified by Ni-NTA affinity chromatography were further purified on SEC. ~40% (37.5% (6/16), 43% (14/32) and 33% (4/12) success rates for the first, second and third approach respectively) had appropriate monodisperse SEC chromatograms that matched the expected theoretical elution profile for the oligomerization state (**fig. 2D**, **fig. S1-2**). These designs were then further examined using either SAXS, negative stain electron microscopy, cryoEM or a combination of techniques (**fig. S1-2)**. Details of the methods, as well as scripts for carrying out the design calculations, are provided in the supplementary materials.

The first approach generated D2, D3 and D4 axle-like structures with folds featuring interdigitated helices with extended hydrogen bond networks. We obtained a 4.2Å 3D reconstruction of a D3 axle (1552_1na0C3_int2_11) which showed close agreement with the design model topology. While the backbone was nearly identical to the design model, the side-chains could be partially elucidated (**fig. 2B, fig. S1, fig. S4, fig. S6)**. SAXS data also showed overall good agreement with the design (**fig. S1**). SAXS and SEC revealed that the middle homohexameric 50 residues long single helices (without appended DHR *wheel* arms) could be solubly expressed and self-assembled into the correct oligomeric state (DSSR2_1552) (**fig. S1**). Another D3 design consisting of 36 residue long single helices was produced via chemical peptide synthesis and assembled into a homohexamer (DSS_310_117, **fig. S1, fig. S7**), while its fusion to C3 wheels generated a bigger D3 oligomer as designed (1na0C3_DSS310_20, **fig. S1**). A D4 peptide homo-oligomer designed using the same approach (D4_1550_700) had SEC and SAXS spectra indicating the designed correct oligomeric state (**fig. S1**). Negative stain EM of a D2 design (D2_1119_7_tj81C2_V39_6) yielded a low resolution 3D reconstruction with the overall features of the design model (**fig. 2D, fig. S1**). The corresponding central 50 residue D2 peptide (D2_1119_7) again expressed solubly and could be purified in the correct oligomeric state (**fig. S1**).

The second approach generated D3, D4, D5 and D8 axle-like structures with folds featuring interdigitated helices with internal cavities for D5 and D8 (in these cases each central helix only forms contacts with two neighboring ones) (**fig. 2B**). We obtained a ~5.9Å electron density map of a D8 design (D8A_1615) revealing a backbone structure nearly identical to the design model (**fig. 2B, fig. S1, fig. S4-5**). This cylinder-shaped homodecahexamer has a previously unobserved fold, with a large central cavity with an end-to-end pore-like feature, contains a nearly straight helix spanning 84 residues and has opposing N and C termini close to its center (**fig. 2B, fig. S1**). Negative stain EM on additional designs: two D8s (D8A_6043 and D8_6_49), one D5 (D5_57C) and one D4 (DC4G1_178), yielded low resolution 3D reconstructions with the features of the design models (**fig. 2D, fig. S1-2**). We converted several of these designs from dihedral to cyclic symmetry by connecting N and C termini, and two such designs, one C5 (C5_41) and one C8 (C8D8_6_49), yielded EM reconstructions with good agreement with the design model (**fig. 2D, fig. S1-2**). Other designs (six D3s, two D4s and one D5s) for which EM data was not obtained were characterized by SAXS and showed similar profiles, which were consistent with the correct oligomeric state and overall structural features (**fig. 2D, fig. S1-2**).

The third approach yielded four C3 axles with folds of smaller aspect ratio and overall size, containing a large wheel-like DHR feature at one end, a narrow central three helix section and a six helix section at the other end. In all cases, the SAXS profiles together with SEC traces suggested that the correct oligomerization state was realized in solution. For design A15.5 we obtained a low resolution cryoEM map that recapitulated the general features of the design model, with prominent C3 symmetric DHR extremities and opposing prism-like extensions (**fig. 2C**, **fig. S2-3**).

## Design of axle-ring assemblies

We next sought to assemble diverse axle-ring assemblies to explore the correspondence between the symmetry and energy landscape of the interface and the mechanical properties. The first challenge was to direct the self-assembly in solution of the ring around the axle by designing energetically favorable interactions, while maintaining some rotational freedom. We first sought to do this by designing assemblies with low residue interaction specificity, loose interface packing, as well as non-obligatory symmetry mismatched interactions between axle and ring restricting only parts of the assembly to form tight contacts (*i.e.* the full interface is never fully satisfied). To achieve these properties, we initially focused on electrostatic interactions between ring and axle which are longer range and less dependent on shape matching than the hydrophobic interactions generally utilized in protein design. To prevent potential disassembly at low concentrations, we aimed to kinetically trap the ring around the axle by installing disulfide bonds at the ring subunit-subunit interfaces. Further, to gain stepwise control on the *in vitro* assembly process, we introduced buried histidine mediated hydrogen bond networks at the ring asymmetric unit interfaces to enable pH controlled ring assembly (**fig. 3A,** see “Experimental methods” in the supplementary materials).

**Fig. 3:**
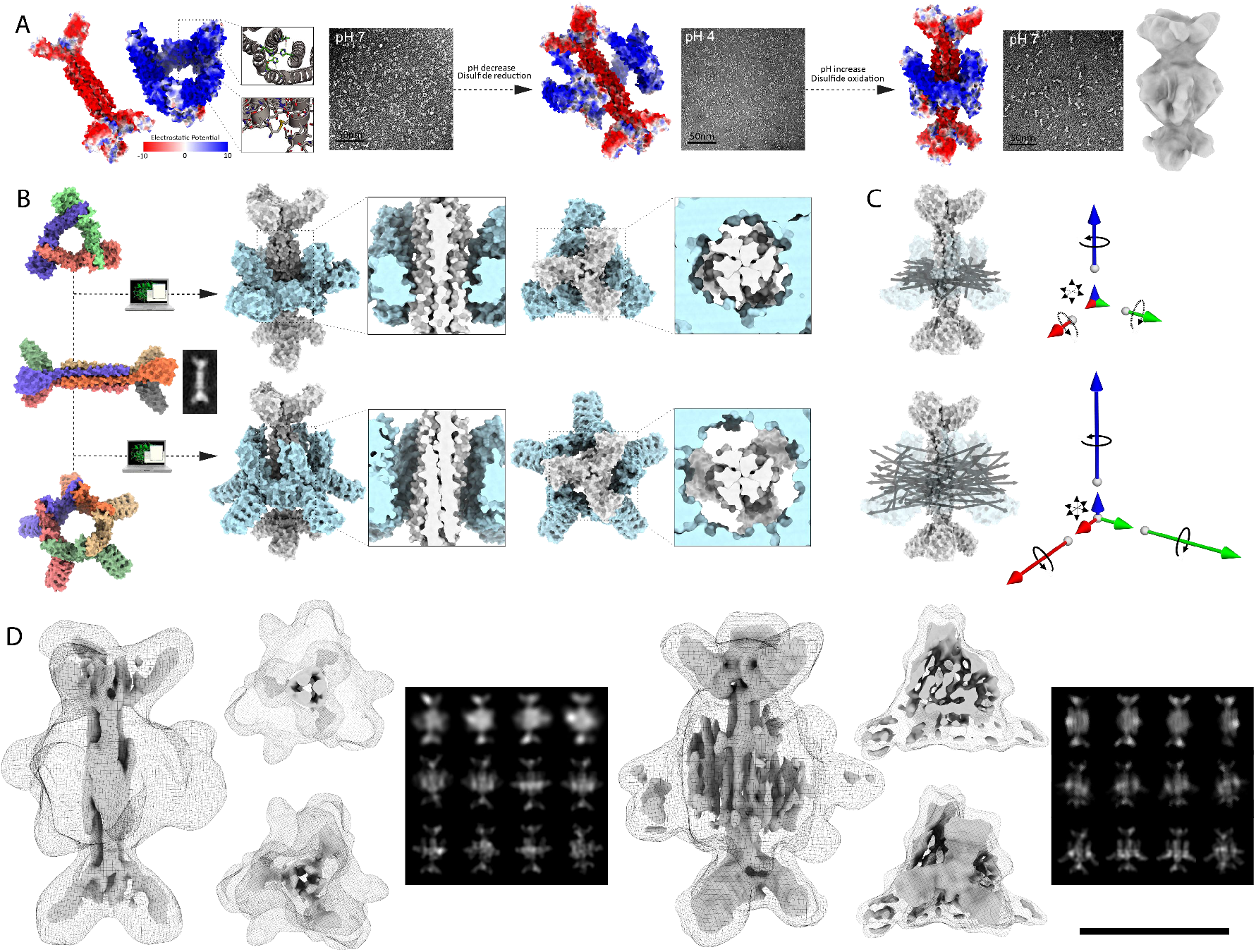
Design of symmetry mismatched D3-C3 and D3-C5 axle-ring assemblies. (**A**) Quasisymmetric axle and ring complex directed self-assembly strategy. Axles and rings are designed with complementary charged residues at their interfaces (electrostatic potential rendered from red to blue), buried histidine bond networks and disulfide bonds across the ring asymmetric unit interfaces to allow pH controlled assembly and oxidoreductive locking of the ring around the axle. Assembly monitored by negative stain EM (square panels) yields fully assembled rotors (cryoEM electron density on right). (**B**) Models of assemblies generated from a D3 axle (1552_1na0C3_int2_11) and C3 (R113) or C5 (C2arms9) rings, and cryoEM 2D average of axle alone before assembly. (**C**) Interface shape and symmetry results in different DOFs. In MD simulations, the D3-C3 system is largely constrained to rotation along the z axis (blue), while the D3-C5 assembly allows rotation along x (green), y (red) and z, and translation in z, x and y. (Left) N-C termini unit vectors of an ensemble of MD trajectories (Right) Vector magnitude corresponding to the computed mean square displacement of the ring relative to the axle along the 6 DOFs. (**D**) 3D CryoEM reconstruction of D3-C3 (Left) and D3-C5 (Right) rotors (axle as surface and ring as mesh, processed in D3 for D3-C3; processed in C1 and shown as surface and mesh at different thresholds for D3-C5; maps are shown as side view, end-on views and transverse slices) and experimental (top row) and theoretical 2D class averages with (middle row) and without (bottom row) explicitly sampling along DOFs. The D3-C3 rotor electron density at 10.2Å resolution suggests that the ring sits midway across the D3 axle consistent with the designed mechanical DOF. The D3-C5 rotor cryoEM electron density at 11.4Å captures the features of the designed structure also evident in the class average (Right). The 2D averages capture secondary structure corresponding to the C5 ring but could not be fully resolved, consistent with the ring populating multiple rotational states. Scale bar for cryoEM density: 10nm

We tested this approach by selecting three of the machine components described above --a D3 axle, a C3 ring and a C5 ring -- and constructing ring-axle rotary machine assemblies with D3-C3 and D3-C5 symmetries (design A113_C2ams9 and C3D3_AR113 respectively, **fig. 3B, fig. S8**). Using PyRosetta(*27*), we threaded axles and rings together by sampling rotational and translational DOF, and designed complementary electrostatic interacting surfaces excluding positively charged residue identities on the axle (Lysine and Arginine) and negatively charged residues (Aspartate and Glutamate) on the ring. Due to the shape complementarity between the internal diameter of the rings and the axle thickness, the interface is tighter for the D3-C3, constraining the ring midway on the axle, and loose for the D3-C5 where the ring can diffuse along multiple DOF, thus resulting in different mechanical constraints: the D3-C3 is only allowed to rotate along the main symmetry axis, while the D3-C5 ring can rotate along x, y and z, as well as translate in z and y (**fig. 3B-C, fig. S15**). Synthetic genes encoding one axle and 2 ring designs were obtained and the proteins were separately expressed in *E. coli* and purified by Ni-NTA affinity chromatography and SEC, which indicated that the surface redesign did not affect the solubility or homo-oligomerization process (**fig. S1-2**). Following stoichiometric mixing of the designed D3 axle and C3 ring, EM analysis showed a collection of assembled and isolated axle and ring molecules (**fig. 3A, left panel**). After dropping the pH and reducing the disulfide, the particles appeared as a mixture of opened, linear and hard to distinguish particles (**fig. 3A, middle panel**). After restoring the pH under oxidizing conditions, the particles appeared fully assembled by EM (**fig. 3A, right panel**). Using biolayer interferometry assays we found that the ring and axle associated rapidly with a Kd in the micromolar range (**fig. S9**). Similar results were obtained with D3-C5 rotary assemblies, and SEC profiles and SAXS spectra were in agreement with the design model in both cases (**fig. S8**).

We next experimented with the design of shape complementary axle and ring components, reasoning that this would enable more precise control of the rotational energy landscape by leveraging Rosetta’s ability to design tightly packed interfaces and hydrogen-bond networks mediated specificity(*25*). We designed four axle-ring assemblies using this approach: a fully C3 symmetric assembly consisting of a C3 axle and a C3 ring (C3-C3, A15.5R82), a symmetry mismatched assembly consisting of a D8 axle around which two C4 rings are assembled (D8-C4, 119RC4_20), a symmetry mismatched rotor consisting of C5 axle and C3 ring (C5-C3_2412 and C5C3_3250), as well as a C8-C4 rotor corresponding to a circular permutation version of the D8-C4 (C8D8_6_49_119RC4_20) (**fig. 4A, fig 4B, fig. S8**). The symmetry matching of the ring and axle in the C3-C3 rotor differs from the mismatching in other assemblies, and the two ring D8-C4 assembly tests the incorporation of multiple coupled rotational DOF in a multicomponent system and also provides a simple way to monitor the position of rings relative to each other by experimental structural characterization, thus providing an indirect way to monitor rotation. Similarly, the DHR arms on other rotors offer direct structurally accessible monitoring of the rotation by visualizing the alignment of axle and ring arms relative to each other. These designs were generated by systematically sampling rotational and translational DOF, removing arrangements with backbone to backbone clashes (**fig. 2B**, see “Computational design methods” in the supplementary materials), and then using the Rosetta HBnet protocol and FastDesign(*28*) to optimize the interface energy. Each interface design trajectory generates widely different periodic energy landscapes according to interface metrics and design specifications (**fig. S10**). In the case of the D8-C4, C5-C3 and C8-C4 designs, since the symmetry of the ring is internally mismatched to the axle, we used a quasisymmetric design protocol (see “Computational design methods” in the supplementary materials). The C4 ring, which is internally C24 symmetric due to the repeated nature of sequences from which it is built, can accommodate the symmetry of D8 or C8 axles since 24 is a multiple of 8, which allows pairing of interactions at the interface while maintaining overall C4 symmetry. In contrast, the C5-C3 arrangement has broken symmetry with a resulting energy landscape with 15 energy minima, with periodicities reflecting the constituent C5 and C3 symmetries (**fig. S10**). This design approach generated shape complementary axle-ring interfaces with an overall cogwheel topology.

**Fig. 4:**
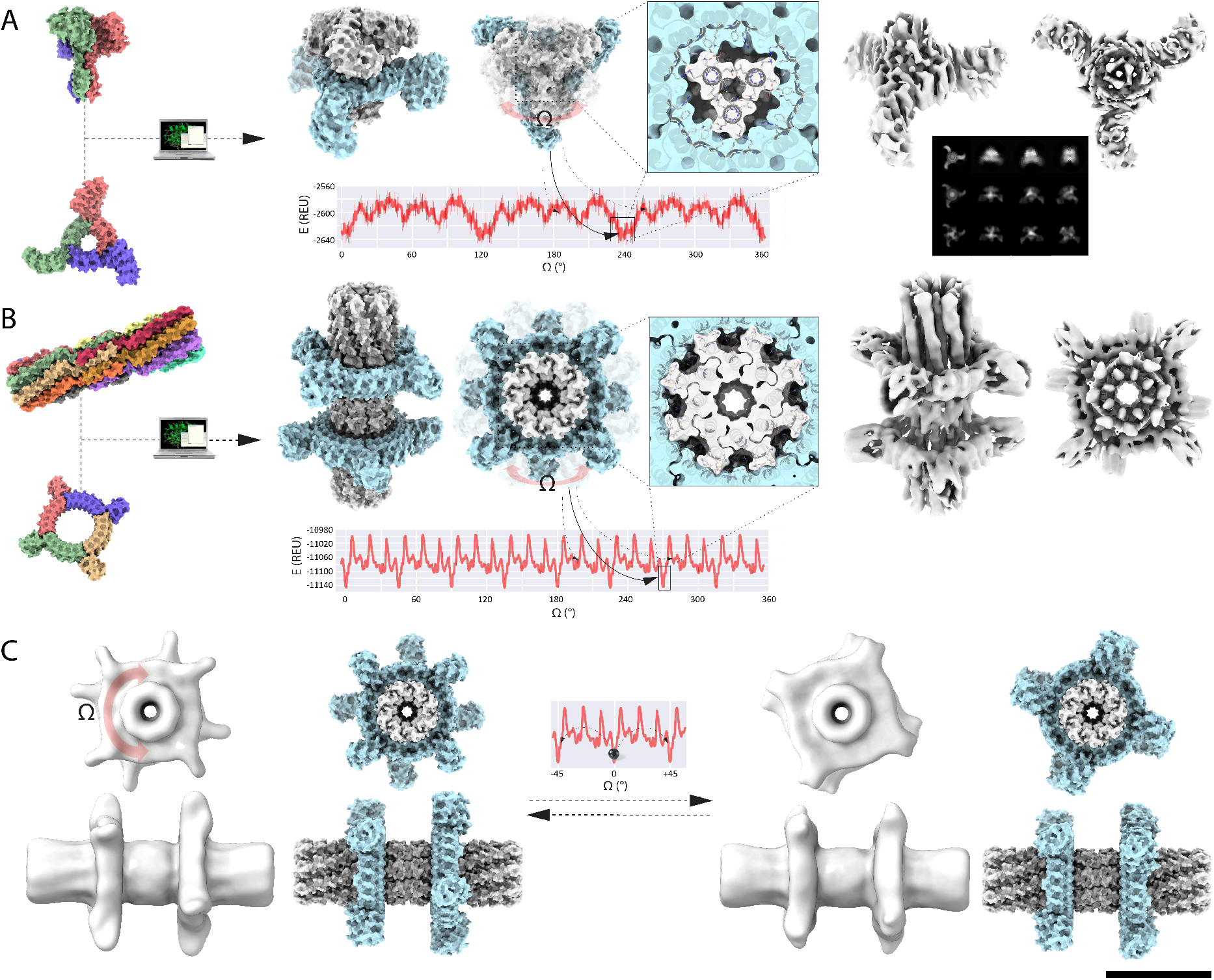
Computational sculpting of the rotational energy landscape by design of interface side-chain interactions. (**A**) Symmetry matched C3-C3 axle and ring complex (Left) Axle, ring, and rotor assembly models. The rotational energy landscape computed by scoring 10 independent Rosetta backbone and side-chains relax and minimization trajectories (solid red line with error bars depicting the standard deviation) features three main energy minima corresponding to the C3 symmetry of the interface with 9 additional lesser energy minima. (Right) Single particle cryoEM analysis of the designed C3-C3 rotor. The electron density (in grey) at 6.5Å resolution shows the main features of the designed structure, evident in the experimental 2D class average (top row) compared to theoretical 2D class averages with (middle row) and without (bottom row) explicitly sampling the DOFs (**B**) Quasisimmetric D8-C4 axle and ring complex (Left) Axle, ring, and rotor assembly models. The rotational energy landscape computed as described in A features eight main energy minima corresponding to the C8 symmetry of the interface (Right) Single particle cryoEM analysis of the designed D8-C4 rotor. The electron density (in grey) at ~5.9Å resolution shows the main features of the designed structure. (**C**) 3D variability analysis of the cryoEM data in relation with the rotational landscape of the D8-C4 rotary machine. The two distinctly resolved structures (shown in light grey) are separated by a 45° rotational step. Scale bar: 10nm

Designs with each of the four symmetries were screened for assembly by expressing ring and axle pairs bicistronically and carrying out Ni-NTA purifications relying on a single HIS tag on the ring component (**fig. S11A**). ~50% (6/12) of C3-C3 designs appeared to express solubly and could be pulled down by the purification process, suggesting that the two components assembled in cells (**fig. S11B**), and one design (54.7.112, **fig. S8**) was further selected for further characterization. The SEC profile in combination with native mass spectrometry indicated an oligomeric state corresponding to the designed assembly, and SAXS data collected on the protein showed good agreement with the design model (**fig. S8, fig. S11C-D**). Using biolayer interferometry we analysed the capacity of the designed axle and ring to assemble *in vitro* into the full rotor, and found that this system showed rapid assembly kinetics with a Kd in the micromolar range (**fig. S9**). Twelve D8-C4 designs were likewise screened for *in vitro* assembly by isolating axle and rings individually by Ni-NTA purifications, and then assayed for assembly by mixing components in stoechiometric fashion. These mixtures were then further SEC purified and the oligomeric assembly state could thus be assessed in addition to SAXS validation, indicating that some of these rotors could self-assemble *in vitro*, while EM data indicated that the rotors were assembling as designed (**fig. S8**). Two out of twelve C5-C3 and one out of six C8-C4 designs tested likewise assembled into axle-ring systems based on SEC chromatograms, and SAXS data, biolayer interferometry binding kinetics and negative stain EM data were consistent with assembly (**fig. S8**, **fig. S9**).

## Population of multiple rotational states

To map the rotational landscape at the single molecule level, we subjected one design from each symmetry class to single particle cryoEM examination. For D3-C3 and D3-C5, we obtained 2D class averages from the collected data that clearly resembled predicted projection maps, and 3D reconstructions in close agreement with the overall design model topology and designed hetero-oligomeric state (**fig. 3D, fig. S12-13, Table S1**). For both designs, the D3 axle was clearly visible and we obtained a high resolution structure nearly identical to the design model. We were able to obtain a high resolution 3D reconstruction map for the D3-C3 rotor assembly, which showed a clear density of the ring sitting in the middle of the axle and recapitulating the C3 ring arms extension, either after processing in C1, C3 or D3 mode (**fig. S12)**. The ring of the D3-C5 design also showed clear density but its resolution could not be further improved as the secondary structure placement relative to the axle were variable, likely due to motion of ring and axle along the multiple DOFs (**fig. S13)**. Cryosparc 3D variability analysis(*29*) suggested that the helical features corresponding to the ring can populate variable positions around the axle according to rotational DOFs only for D3-C3, and translational and rotational DOFs for D3-C5 (**fig. 3B-C**, **Movie S1-4**). This is also evident from visual inspection of the cryoEM 3D reconstruction: the ring arms populate multiple positions along the rotational axis (**fig. 3D**). Explicit modelling of rotational variability along the designed DOFs was necessary to produce theoretical projections closely resembling the experimental 2D class averages (**fig. 3D, fig. S14**). Molecular dynamics simulations (MD) recapitulated the intended internal rotary motion between ring and axle, with the D3-C5 rotary machine showing increased displacement along allowed DOFs compared to D3-C3 (**fig. 3C**, **fig. S15**). Taken together, the cryoEM data and molecular dynamics simulations are consistent with the design goal of constrained internal rotation.

Single particle cryoEM analysis of a C3-C3 assembly yielded 2D class averages with the axle and ring clearly visible. We were able to generate a 3D reconstruction with a resolution of 6.5Å, which yielded an electron density map similar to the design model (**fig. 4A, fig. S3, fig. S8, Table S1**). However, the high orientation bias of the particle in ice considerably limited the resolution of the structure by preventing the obtention of side views. We hypothesize that the diffuse density of the axle in the middle of the clear ring in top view class averages could be attributed to rotational diffusion (**fig. 4A, fig. S3**). This appeared evident after explicitly modeling rotational variability along the designed DOF, which produced theoretical averages closely resembling the experimental data (**fig. 4A, fig. S14**). This is consistent with the designed smooth energy landscape with 3 energy minima at a 60° rotation distance and 9 other 30° spaced degenerate alternative wells separated by energy barriers.

The predicted energy landscape of the D8-C4 design is quite rugged, with a total amplitude of 151.7 REU with 8 steep wells spaced 45° stepwise along the rotational axis corresponding to the high symmetry of the interface. We obtained a cryoEM map of ~5.9Å resolution very close to the design model (**fig. 4A, fig. S5, Table S1**). 3D variability analysis calculations using Cryosparc software(30) showed that the experimental structural data could be clustered in two nearly equiprobable states which corresponded to two rotational states of one ring relative to the other, corresponding to pronounced energy-minima with 45° steps along the rotational axis consistent with the *in silico* designed energy landscape. There are two clearly identifiable structures in which the ring arms are either aligned or offset, as in the eclipsed and staggered arrangements of ethane (**fig. 4C, fig. S5, Movie S5**). While cryoEM provides a frozen snapshot of rotational bins, this data shows that the system can assemble and sample mechanical rotational bins according to the design specifications. Taken together, these results suggest that the explicit side-chain interaction design reduces the degeneracy of rotational states observed with purely electrostatic interactions.

## Conclusions

Our proof of concept rotary machine assemblies demonstrate that protein nanostructures with internal mechanical constraints can now be designed. The hetero-oligomers topologies we created do not exist in nature nor have such synthetic systems been designed previously, and provide insights towards the design of more complex protein nanomachines. First, systematic and accurate *de novo* design according to machine components specification (**fig. 1**, **fig. 2**), coupled with computational sculpting of the interface between parts can be used to simultaneously promote self-assembly and constrain motion along internal degrees of freedom. Second, the shape and periodicity of the resulting rotational energy landscape is determined by the symmetry of components, the shape complementarity of the interface, and the balance between hydrophobic packing and conformationally promiscuous electrostatic interactions (**fig. 3**, **fig. 4A-C**). Symmetry mismatch tend to generate assemblies with larger numbers of rotational energy minima than symmetry matched ones, and explicit design of close sidechain packing across the interface results in deeper minima and higher barriers than non-specific interactions (**fig. 3, fig. 4, fig. S10**). In general, the surface area of the interface between axle and ring scales with the number of subunits in the symmetry, resulting in a larger energetic dynamic range accessible for design (**fig. S10**). The combination of the structural variability apparent in the cryoEM data of D3-C3, D3-C5 and C3-C3 designs (**fig. 3D, fig. 4, fig. S3, fig. S12-14)**, the MD simulations (**fig. 3C**, **fig. S15**), and the discrete states observed for the D8-C4 design (**fig. 4C, fig. S5**), suggests that these assemblies sample multiple rotational states. Time-resolved characterization of the internal motion at the single molecule level will reveal how the ability to computationally shape rotational energy landscapes can be used to control Brownian dynamics.

The internal periodic but asymmetric rotational energy landscapes of our designed rotary machine assemblies provide one of two needed elements for a directional motor. An energy harvesting process to break detailed balance and transfer the system into an excited state remains to be designed: for example the interface between machine components can be designed for binding and catalysis of small molecule fuels(*19*). Symmetry mismatch, which plays a crucial role in torque generation in natural motors(*31–37*), can be leveraged for the design of synthetic protein motors. Modular assembly could lead to compound machines for advanced operation or integration within nanomaterials. In this direction, we recently designed modular rotor complexes with reversible heterodimer extensions binding components of the rotor (**fig. S16**). Our protein nanomachines can be genetically encoded for multicomponent self-assembly within cells (**fig. S11**) or *in vitro* (**fig. S9**), facilitating fabrication or *in vivo* transfer and use. Taken together, these approaches could ultimately enable the engineering of a vast range of nanodevices for medicine, material sciences or industrial bioprocesses. More fundamentally, *de novo* design provides a bottom-up platform to explore the critical principles and mechanisms underlying nanomachine function that complements long standing more descriptive studies of the elaborate molecular machines produced by natural evolution.

## Acknowledgements

We thank Bridget Carragher, Clint Potter, Ed Eng, Laura Yen, and Misha Kopylov of the New York Structural Biology Center, for assistance and helpful discussions. We especially thank Sjors Scheres of MRC-LMB for helpful discussions and guidance regarding cryoEM data processing. We would like to thank the Rosetta@Home user base for donating their computational hours to run forward folding simulations. We thank Florian Busch and Vicki Wysocki at Ohio State University for providing expert support with native mass spectrometry experiments and Vicki Wysocki at Ohio State University for providing expert support with native mass spectrometry experiments. We especially thank Danny Sahtoe for all the scientific support and many insightful discussions. Thanks to Florian Praetorius for the brainstorming sessions dedicated to designing *de novo* protein motors. An additional thanks to Asim Bera and Matt Bick, and Justin Decarreaux for support in crystallography and optical microscopy respectively. Thanks to Tom Daniel for all the great interactions and fascinating ideas and discussion and Luis Ceze for accompanying the thought process regarding the design of protein nanomachines, either computational and mechanical. Thanks to Brian Coventry for very helpful advice and computational help, and Lance Stewart for expert help, advice, perspectives and discussions. Finally, special thanks to Ashley Nord for invaluable discussions and expert insights into the complexity of natural molecular motors biophysics.

## Funding

National Science Foundation (NSF) award 1629214 (DB)

A generous gift from the Audacious Project (DB)

the Open Philanthropy Project Improving Protein Design Fund (DB)

University of Washington Arnold and Mabel Beckman cryo-EM center (DB, DV, JK, JQ)

A S10 award funded the purchase of a Glacios microscope, award number S10OD032290 (DB, DV, JK, JQ)

NIH Molecular Biology Training Grant T32GM008268 (YH)

Human Frontiers Science Program Long Term Fellowship (AC)

Washington Research Foundation Senior Fellow (AC)

Howard Hughes Medical Institute research (AC)

SAXS data was collected at the Advanced Light Source (ALS), SIBYLS beamline on behalf of US DOE-BER, through the Integrated Diffraction Analysis Technologies (IDAT) program. Additional support comes from the NIGMS project ALS-ENABLE (P30 GM124169) and a High-End Instrumentation Grant S10OD018483. The Berkeley Center for Structural Biology is supported in part by the National Institutes of Health (NIH), National Institute of General Medical Sciences, and the Howard Hughes Medical Institute.

The Advanced Light Source (ALS) is supported by the Director, Office of Science, Office of Basic Energy Sciences and US Department of Energy under contract number DE-AC02-05CH11231.

Some of this work was performed at the Pacific Northwest Center for Cryo-EM, which was supported by NIH grant U24GM129547 and performed at the PNCC at OHSU and accessed through EMSL (grid.436923.9), a DOE Office of Science User Facility sponsored by the Office of Biological and Environmental Research.

Molecular graphics and analyses were performed with UCSF Chimera, developed by the Resource for Biocomputing, Visualization, and Informatics at the University of California, San Francisco, with support from NIH P41-GM103311.

## Author contributions

Conceptualization: AC, DB

Methodology: AC, DB, JH, JK

Software: AC, YH, CX, SB, GU, UN, PB, DB, DS, AM, NK, WS, NB

Validation: AC, DB, JH, JK, YH

Formal analysis: AC, JH, NB

Investigation: AC, JH, NB, YJ, DN, JQ

Resources: AC, DB, JH, JK, JQ

Data curation: AC, JH, YH

Writing – original draft: AC

Writing – review & editing: AC, DB, JH, YH, JK

Visualization: AC, JH, NB

Supervision: DB, JK

Project administration: AC, DB

Funding acquisition: DB, JK, DV, AC, YH

## Competing interests

Authors declare that they have no competing interests.

Data and materials availability

All data are available in the main text or the supplementary materials.

## Supplementary Materials

Materials and Methods

Figs. S1 to S16

Tables S1

Movies S1 to S5

Data S1 to S5

## Notes

### Competing Interest Statement

The authors have declared no competing interest.

