## Supplementary materials and methods for "Computational design of nanoscale rotational mechanics in *de novo* protein assemblies"

**This PDF file includes:**

Materials and Methods
Figs. S1 to S16
Tables S1
Captions for Movies S1 to S5

**Other Supplementary Materials for this manuscript include the following:**

Movies S1 to S5
Data S1 to S5

All data is accessible on our server at https://files.ipd.uw.edu/acourbet/Data_S*zip

### Materials and Methods

Computational Design

Generation of homooligomeric dihedral and cyclic symmetric axle parts:

Approach 1: This approach relies first on the design of short (30 to 50 residues) single alpha helices monomers self-assembling into high aspect ratio dihedral homoligomers, which are then further fused to cyclic wheel shaped homooligomers to yield full axle parts.

Parametric design was used to generate short single α-helices and sample backbone configurations by systematically varying helical parameters using the Crick generating equations(*24,38*). As described before, ideal values were used for the supercoil twist (ω0) and helical twist (ω1).

In the case of D3 helical bundles, in order to obtain the helical interdigitated geometry allowing the obtain a packed core without holes after design and therefore assembly of single helices into dihedral symmetry, we sampled two segments with different starting point for the superhelical radii per helix (6Å and 12Å), joined by a custom number of linker residues between the two segments, using a custom python script. This parameter range (~5Å with bins of 0.5Å) were chosen based on iteratives cycles of parametric helix generation and Rosetta design with metric assessment, and the range of metrics yielding the highest scoring backbone were chosen. The helical phase (Δϕ1) was sampled from 0° to 90° with a step size of 10°. We sampled the offset along the z-axis (Z-offset) from -1.51Å to 1.51Å, with a step size of 0.1Å. The supercoil phases (Δϕ0) were fixed at 0°, and 30° for D3s and D2s, respectively.

Once ideal backbones geometry were generated using this parametric approach, we used the Rosetta design protocol to further design side chains identities and rotamers and optimize the interface energy to direct the assembly in dihedral homooligomers. Importantly, this step relied on the use of the Rosetta HBnet protocol described previously(*25*), which allows for extended hydrogen bond networks across monomer subunits therefore ensuring specificity of interaction and symmetric binding mode.

The dihedral building blocks were then rigidly fused to previously designed cyclic homooligomers(*39*) by designing short rigid helical linkers bridging the two building blocks. The inner helices of the dihedral assemblies obtained (C or N termini depending on design) were then fused by short structured helical fragments using Rosetta Remodel(*40*) while sampling the rotation and distance between Z aligned cyclic homooligomers and dihedral homooligomer. To further stabilize and optimize the generated Cyclic-Dihedral fusion, a second round of Rosetta design of the fusion was performed.

Method 2: This approach relied on alpha helical extensions of N or C termini of previously designed cyclic homooligomers, in order to direct the assembly of two elongated cyclic homooligomers into high aspect ratio dihedral symmetric axle parts. Rosetta SymDofMover was used to set up the symmetry in which the input monomer subunits were aligned along the z axis. Input subunits were first optionally flipped 180 degrees about the z axis to reverse the inputs if necessary, so that the N or C termini to be elongated would point toward each other. Monomer subunits were then translated along the specified z axis and rotated about the z axis according to random Gaussian sampling in order to finely sample helical extension parameters. Following these initial manipulations of the input structures, a symmetric pose was generated using D3, D4, D5, D6 or D8 symmetry definition files. We then applied the Rosetta BluePrintBDR mover which allowed us to build helical fragment extension starting at the previously positioned monomers, and spanning the distance between symmetric subunits. Once centroid helical backbones geometries were generated and sampled, we used the Rosetta design protocol to further design side chains identities and rotamers and optimize the interface energy to direct the assembly in dihedral homooligomers. Importantly, this step relied on the use of the Rosetta HBnet protocol described previously(*25*), which allows for extended hydrogen bond networks across monomer subunits therefore ensuring specificity of interaction and symmetric binding mode.

Generation of cyclic symmetric homooligomeric rotor parts:

Computationally designed ring shape structures or various symmetries (C1, C3, C4), were either collected from previously published work(*21,41*), or designed from heterodimers and DHRs in symmetry mode (C3, C5) using protocols previously described(*12*). 9x, 12x and 24x toroids were used in C1 symmetric versions or cut into 3 or 4 to produce C3 or C4 symmetric homooligomers. All designs were then computationally augmented by systematic symmetric fusion of DHR repeats proteins using the HFuse protocol, and the surrounding fusion interface of the fusion was further redesigned using Rosetta design protocols to optimize the assembly energy.

Generation of two component rotary machine models from symmetric axle and rotor parts:

The goal of the computational docking procedure between axle and rotor machine parts was to exhaustively sample the rotational conformational space within some specified resolution and meaningful interface quality, all possible ways to assemble a full rotary machine complex from the two libraries of previously designed axle and rotor parts.

We started by enumerating all possible rotary machine assemblies by inspecting shapes and dimensions of available parts and identifying assemblies that would not produce any steric clashes. We then proceeded to computational docking of parts using a two-dimensional rigid body docking space to allow contact between the axle and rotor (one rotation and one translation along the Z axis). We sampled 180° rotation for C2s, 120° for C3s, 90° for C4s, 72° for C5s, and we sampled the whole span on available translation along the axis that would not generate clashes between backbones, with a 1° and 1A step, respectively. For each sampled dock, the resulting heteromultimeric interface was designed either using Rosetta design and HBnet to obtain tightly packed, specific interfaces with extended hydrogen bond networks, and in some cases by constraining the residue identities of the axle (DEHQTNSY) and ring (KRHQTNSY) to obtain complementary charges allowing loose non specific interactions. Since some of the resulting assemblies have intrinsic symmetry mismatch between the axle and rotor (*e.g*. D8 axle and C4 ring), we used a quasi-symmetric design methodology, relying on the Rosetta StoreQuasiSymmetricTaskMover, which creates a stored task that links selected interface residues. The residues remain identical in identity when the interface is designed, but their rotamers are packed differently, which allows identical residues in symmetric subunits to satisfy multiple interfaces at the same time.

In order to kinetically trap rings onto the axle, we further generated a disulfided version of homooligomers by placing cysteine at the interface between asymmetric units. This was achieved using a PyRosetta based stapling method that allows to identify pairs of residues that can accommodate disulfides given the 3D structure of a protein (https://github.com/atom-moyer/stapler).

Interface disulfide stapling

This protocol was developed to quickly identify pairs of residues that can accommodate disulfides given the 3D structure of a protein. 30,000 native disulfide structures were procured from the PDB, and the relative positions of the backbone atoms (N, CA, C) were calculated, hashed, and stored into a database. A candidate protein structure can then be searched for residue pairs at all relative positions of backbone atoms that can accommodate disulfides according to native geometries.
(https://github.com/atom-moyer/stapler)

Molecular dynamics simulations

Rosetta models of D3-C3 and D3-C5 with truncated ring DHR arms (to minimize the total number of atoms to simulate) were used as the starting coordinates for the simulations. The rotor rings of D3-C3 and D3-C5 were rotated at 10 and 12 degree intervals, respectively. Each model was solvated in an octahedral periodic box of OPC water and 70 mM NaCl using AmberTools18(*42*). In total, each system consisted of approximately 590,000 atoms. Simulations were run at constant pressure (1 bar) and temperature (298 K) using the Monte Carlo barostat, the Langevin thermostat and the ff19SB forcefield(*43*). Using the CUDA enabled version of Amber18, four parallel simulations for each rotated model were equilibrated using the AmberMDprep protocol(*44*). Once equilibrated, the simulations were run at 2 fs timestep for a total of 40 ns each, yielding an aggregate simulation time of 1920 ns for D3-C3 and 960 ns for D3-C5. To allow exploration of the rotors' degrees of freedom from the initial configurations, the first 20 ns of each simulation was discarded and the final 20 ns was used in later analysis. To investigate the movement of the rings around their respective axles, 200 ps snapshots of the simulations were aligned to the initial axle coordinates by rmsd. Number density maps of the backbone atoms were calculated using the VolMap command in AmberTool's cpptraj(*45*). These maps were contoured to 0.001 as shown in **fig. S15**. To calculate the axle drift with respect to ring rotation, the backbone center of mass of the rings was calculated for all aligned snapshots. The snapshots were binned according to the ring rotation in 24 degree intervals and then averaged as shown in **fig. S15**. To calculate ring tilt, the centers of mass of each ring subunit was calculated, then a plane was fit through these points using the least squares optimizer in SciPy(*46*). The angle between this plane and the long axis of the axle was taken as the tilt, and this was averaged over rotation as described for the axial drift. The Mean Square Displacement for the DOFs was computed as MSD=average(r(t)-r(0))^2.

All PDBs can be found in DataS5. All scripts and computational methods are accessible on GitHub (https://github.com/alexiscourbet)

Buffer and media recipe for protein expression

TBM-5052: 1.5% [wt/vol] tryptone, 2.5% [wt/vol] yeast extract, 0.5% [wt/vol] glycerol, 0.05% [wt/vol] D-glucose, 0.2% [wt/vol] D-lactose, 25 mM Na2HPO4, 25 mM KH2PO4, 50 mM NH4Cl, 5 mM Na2SO4, 2 mM MgSO4, 10 µM FeCl3, 4 µM CaCl2, 2 µM MnCl2, 2 µM ZnSO4, 400 nM CoCl2, 400 nM NiCl2, 400 nM CuCl2, 400 nM Na2MoO4, 400 nM Na2SeO3, 400 nM H3BO3

Lysis buffer: 25 mM Tris, 25 mM NaCl, 20 mM Imidazole, pH 8.0 at room temperature

Wash buffer: 25 mM Tris, 25mM NaCl, 20 mM Imidazole, pH 8.0 at room temperature

Elution buffer: 25 mM Tris, 25 mM NaCl, 200 mM Imidazole, 50mM EDTA, pH 8.0 at room temperature

TBS buffer: 25 mM Tris pH 8.0, 25 mM NaCl

Construction of synthetic genes

Prior to transformation and expression in E coli hosts, synthetic genes were ordered either from Integrated DNA Technologies (Coralville, IA) or Genscript Inc. (Piscataway, N.J., USA) and cloned in pET29b+ e. coli expression vector between the NdeI and XhoI sites. For bicistronic constructs used for screening the *in cellulo* assembly of axle and rotors, a synthetic bicistron containing both axle and rotor genes were synthesised and cloned at once in the Nde1/XhoI site, with a termination and strong ribosomal binding site sequence between the genes. For most synthetic gene constructs, a C or N ter hexahistidine tag was added in frame after a short GS linker. A stop codon was introduced at the 3' end of the protein coding sequence to prevent expression of the C-terminal hexahistidine tag in the vector.
(See Data_S4_components_and_machines_sequences.fasta for all sequences)

Protein expression

Plasmids were transformed into chemically competent E. coli expression strain BL21(DE3*) (New England Biolabs) for protein expression. Following transformation and overnight growth on Luria-Bertani agar Kanamycin plates 100ug/ml, single colonies were picked and directly transferred into 2x50 ml TBM-5052 medium containing 150 µg/mL Kanamycin and incubated with shaking at 225 rpm for 24 hours at 37°C following the autoinduction method(*47*). After 24 hours of incubation, the temperature was dropped for an overnight incubation at 20°C before harvesting the cells via centrifugation at 4500G for 20 minutes at 4°C.

Affinity purification

The cell pellets were resuspended in 30ml lysis buffer, followed by cell lysis via sonication at 85% power for 2.5 minutes (10 sec on/10 sec off) while keeping the cell suspension at 4°C. Lysates were clarified by centrifugation at 4°C and 18000 G for 45 minutes and applied to columns containing Ni-NTA (Qiagen) resin pre-equilibrated with lysis buffer. The columns were washed 3 times with 10 column volumes (CV) of wash buffer, followed by 15ml of elution buffer for protein elution.

Size-exclusion chromatography (SEC)
Protein elutions were further concentrated in 15mL 3K protein concentrators (Millipore Sigma) to a volume of 500uL and the buffer exchanged for TBS buffer. The resulting protein solutions were purified by SEC using a Superdex 6 10/300 GL increase column (GE Healthcare) or a Superdex 200 10/300 GL increase column in TBS buffer. SEC elution fractions corresponding to the designs theoretical elution volumes were concentrated in TBS prior to further biochemical analysis. The theoretical SEC elution volumes were computed using the following calibrated equations: $V_{S200} = -1.89\log(<\text{mass of design in kDa}>) + 21.9$ ; and $V_{S6} = -1.33\log(<\text{mass of design in kDa}>) + 21.9$ . All SEC traces can be found in DataS3.

D3-C3 and D3-C5 assembly process
D3 axles and C3 or C5 rings were purified as previously described. Axle and ring were then mixed in TBS solution with 25mM TCEP following a 1:1 stoichiometry, after which the pH is dropped to 3.0 by dialysis in citrate buffer with TCEP. The protein samples were then heated for an hour at 65C, and then allowed to cool back down to room temperature on a bench. The protein samples were then dialysed overnight in TBS buffer and further SEC purified.

Small Angle X-ray Scattering (SAXS)
Protein samples were purified by SEC in 25 mM Tris pH 8.0, 25 mM NaCl and 1% glycerol; elution fractions corresponding to the protein were further concentrated using 3K protein concentrators (Millipore Sigma) and the flow-through was used as blank for buffer subtraction. SAXS Scattering measurements were performed at the SIBYLS 12.3.1 beamline at the Advanced Light Source. The sample-to-detector distance was 1.5 m, and the X-ray wavelength (λ) was 1.27 Å, corresponding to a scattering vector q ($q = 4\pi \sin\theta/\lambda$, where 2θ is the scattering angle) range of 0.01 to 0.3 $Å^{-1}$. A series of exposures were taken of each well, in equal sub-second time slices: 0.3-s exposures for 10 s resulting in 32 frames per sample. For each sample, data were collected for two different concentrations to test for concentration-dependent effects; 'low' concentration samples corresponded to 1 mg/ml and 'high' concentration samples to 5 mg/ml. Collected data were processed using the SAXS FrameSlice online server and analysed using the ScÅtter software package(*23*). The FoXS software (Sali Lab) was used to compare experimental scattering profiles to design models and assess quality of fit(*48-50*). All SAXS traces can be found in DataS2.

Electron microscopy
Negative Stain Electron Microscopy:
SEC fractions corresponding to the designs were concentrated in TBS prior to negative stain EM screening. Samples were then immediately diluted 5 to 150 times in TBS buffer (tris 25mM, NaCl 25mM) depending on the concentration of the samples. A final volume of 5 µL was applied on negatively glow discharged, carbon-coated 400-mesh copper grids (01844-F, TedPella,Inc.), then washed with Milli-Q Water and stained using 0.75% uranyl formate as previously described(*51*). Air-dried grids were then imaged on either a FEI Talos L120C TEM (FEI Thermo Scientific, Hillsboro, OR) equipped with a 4K × 4K Gatan OneView camera at a magnification of 57,000x and pixel size of 2.51Å. Micrographs collection was automated using EPU software (FEI Thermo Scientific, Hillsboro, OR) and were imported into CisTEM software(*52*) or cryoSPARC software(*53*). CTF estimation was done with CTFFIND4 and a circular blob picker was used to select particles which were then subjected to 2D classification.

Ab initio reconstruction and homogeneous refinement in Cn symmetry were used to generate 3D electron density maps.

<u>CryoEM Sample Preparation and DataCollection:</u>

CryoEM grids were prepared by diluting protein samples with TBS 1 to 10 times immediately before applying 3.5 µL to glow-discharged 400 mesh, C-flat, 2 micron holes, 2 micron spacing, CF-2/2-4C (CF-224C-100) (Electron Microscopy Sciences, Hatfield, PA) cryoEM grids. For some samples, multiple blots were applied in order to obtain the best particle density. All grids were blotted using a blot force of 0 and 5 second blot time at 100% humidity and 4°C and plunge-frozen in liquid ethane using a Vitrobot Mark IV (FEI Thermo Scientific, Hillsboro, OR). All cryoEM grids were screened on a Glacios transmission electron microscope (FEI Thermo Scientific, Hillsboro, OR) operated at 200 kV and equipped with a Gatan K2 Summit direct detector. Automated glacios data collection was carried out using Leginon(*54*) at a nominal magnification of 36,000x (1.16 Å/pixel). Movies were acquired in counting mode fractionated in 50 frames of 200 ms at 8.5 e-/pixel/sec for a total dose of ~65e-/Å$^2$. High resolution data was collected on a Titan Krios (FEIco.) operating at 300kV, with a Quantum GIF energy filter (GatanInc.) operating in zero-loss mode with a 20eV slit width, and a K-2 Summit Direct Detect camera. Movies were acquired using Leginon in super-resolution mode at 130,000X (pixel size 0.525Å/pixel) with 50 frames at an exposure rate of 2.5 e-/pixel/sec for a total dose of ~90e/Å$^2$. Details of dataset processing for each design are illustrated in Table S1 and Figure S3, S5, S6, S12 and S13. Theoretical 2D projections were generated using CryoSparc software's "create template" function from an input volume generated with EMAN2(*55*).

<u>CryoEM data processing:</u>

Multiple datasets were collected for each design and combined early on during processing. See table 1 and processing flowcharts for details. Briefly, images were manually curated to remove poor quality acquisitions such as bad ice or large regions of carbon. Dose-weighting and image alignment of all 50 frames was carried out using MotionCor2(*56*) with 5X5 patch or with cryosparc v2 patch alignment tool with default parameters. Super-resolution krios data was binned 2X during alignment. Initial CTF parameters were estimated using CTFfind4(57). Particle picking was done with a gaussian blob picker and in some cases followed by a template picker. Particles were extensively classified in 2D to remove junk particles and designs which may not have been intact or were damaged, yielding in some cases relatively few particles. This may also be due to the low mass of the designed proteins which did not align well. In addition, the expected motion of the rotors may have introduced further heterogeneity, limiting classification efforts. Starting models for all designs were always obtained *ab initio*, despite clear evidence of the expected design in 2D. In 3D classification and refinement we were able to resolve either axle or ring, and in one case both together (D8-C4), suggesting rotor movement. FSC 0.143 curves were generated by exporting half maps to relion for post-process. Local resolution estimates were generated in relion and displayed onto the locally filtered map outputs using Chimera(*58*). For density modification in Phenix(*59*), we used as input the exported half maps from cryosparc with default params at 100 bins and local filtering with a factor of 5. FSC curves were plotted using the Phenix density modification Fref 0.5 output along with the relion FSC estimates. Directional FSC calculated using remote 3DFSC processing tool (https://3dfsc.salk.edu/). 3D Variability analysis (3DVA) of the D8-C4 design was done in cryosparc v2 following expanded particles in D4 symmetry of the final reconstructions with a mask around both rings and the axel. We used default settings of simple cluster mode and 10 frame output with a 10Å lowpass filter for assessing variability. First and last frames of the second trajectory component were used as input for downstream refinement of distinct structures. Resulting maps were then low-pass filtered to 15Å for clarity. For D3-C5, 3DVA was carried out after D3 symmetry was expanded and variability was processed and filtered at 5Å for display. All electron density maps can be found in DataS1.

Biolayer interferometry

Biolayer interferometry experiments were performed on an OctetRED96 BLI system (ForteBio, Menlo Park, CA). Enzymatic protein biotinylation was performed on SEC purified Avi-tagged proteins prior to the assay. The BirA500 (Avidity, LLC) biotinylation kit was used to biotinylate protein from the IMAC elution according to the manufacturer protocol. Reactions were incubated at 4C overnight and purified using size exclusion chromatography on a Superdex 6 10/300 Increase GL (GE Healthcare) in TBS buffer (25 mM Tris pH 8.0, 25 mM NaCl). Streptavidin coated biosensors were equilibrated for 10 minutes in Octet buffer (10 mM HEPES pH 7.4, 25 mM NaCl, 3 mM EDTA, 0.05% Surfactant P20) supplemented with 1 mg/ml Bovine Serum Albumin (SigmaAldrich). Enzymatically biotinylated axle components were immobilized onto the biosensors by dipping the biosensors into a solution with 10-50 nM protein for 200-500s. This was followed by dipping in fresh octet buffer to establish a baseline. Titration experiments were performed at 25 °C while rotating at 1,000 r.p.m. Association of rings rotor components with axle immobilized on the tips was allowed by dipping biosensors in solutions containing designed protein diluted in octet buffer followed by dissociation by dipping the biosensors into fresh buffer solution in order to monitor the dissociation kinetics.

Native Mass Spectrometry

The oligomeric state of *in vivo* assembled rotors was analyzed by online buffer exchange MS(*60*) using a Vanquish UHPLC coupled to a Q Exactive Ultra-High Mass Range (UHMR) mass spectrometer (Thermo Fisher Scientific) modified to allow for surface-induced dissociation (SID) similar to that previously described(*61*). 1 µL of 25 µM protein in TBS buffer were injected and online buffer exchanged into 200 mM ammonium acetate, pH 6.8 by a self-packed buffer exchange column (P6 polyacrylamide gel, Bio-Rad Laboratories) at a flow rate of 100 µL per min. A heated electrospray ionization (HESI) source with a spray voltage of 4 kV was used for ionization. Mass spectra were recorded for 1000 –20000 m/z at 3125 resolution as defined at 400 m/z. The injection time was set to 200 ms. Voltages applied to the transfer optics were optimized to allow for ion transmission while minimizing unintentional ion activation, and a higher-energy collisional dissociation of 5 V was applied. Mass spectra were deconvolved using UniDec V4.2.2 22. Deconvolution settings included mass sampling every 10 Da, smooth charge states distributions, automatic peak width tool, point smooth width of 1 or 10, and beta of 50.

Visualization and figures

All structural images for figures were generated using PyMOL, Chimera or ChimeraX. Data was processed and figures were plotted using Pandas, MatplotLib, and Seaborn python libraries. Figures were further rendered and assembled using Adobe Illustrator.

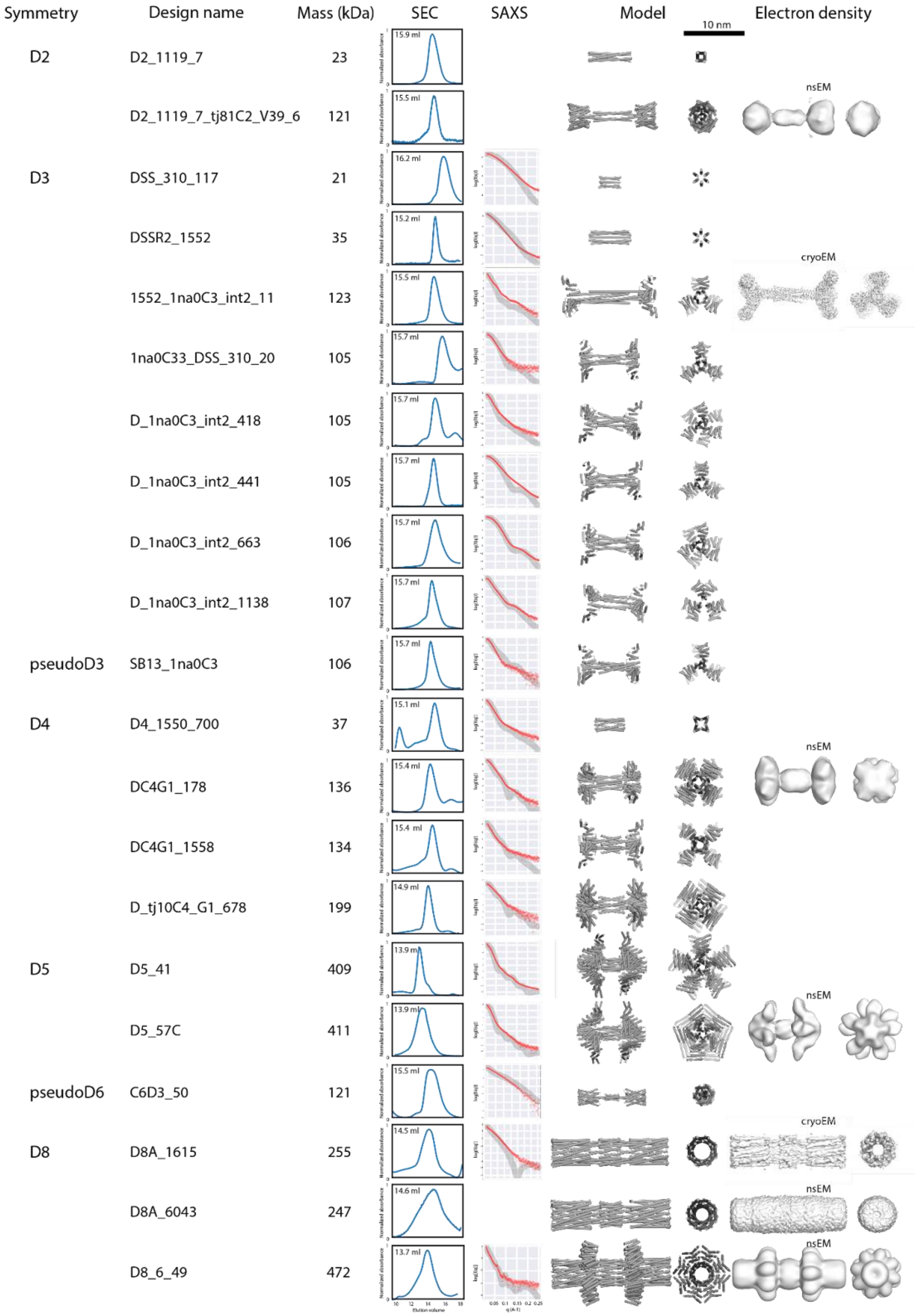


**Figure S1: Detail of the library of axle parts for the design of rotary machines with corresponding symmetry, design nomenclature, oligomeric mass, SEC chromatograms, SAXS traces, designed PDB model, and 3D electron density reconstruction from electron microscopy analysis.** For each SEC trace, the theoretical elution volume corresponding to the correct oligomer state is given in milliliters in black next to the chromatogram. Experimental SAXS traces are shown in the red line, while the theoretical trace corresponding to the design is shown in grey. nsEM: data obtained using negative stain electron microscopy; cryoEM: data obtained using single particle cryoelectron microscopy.

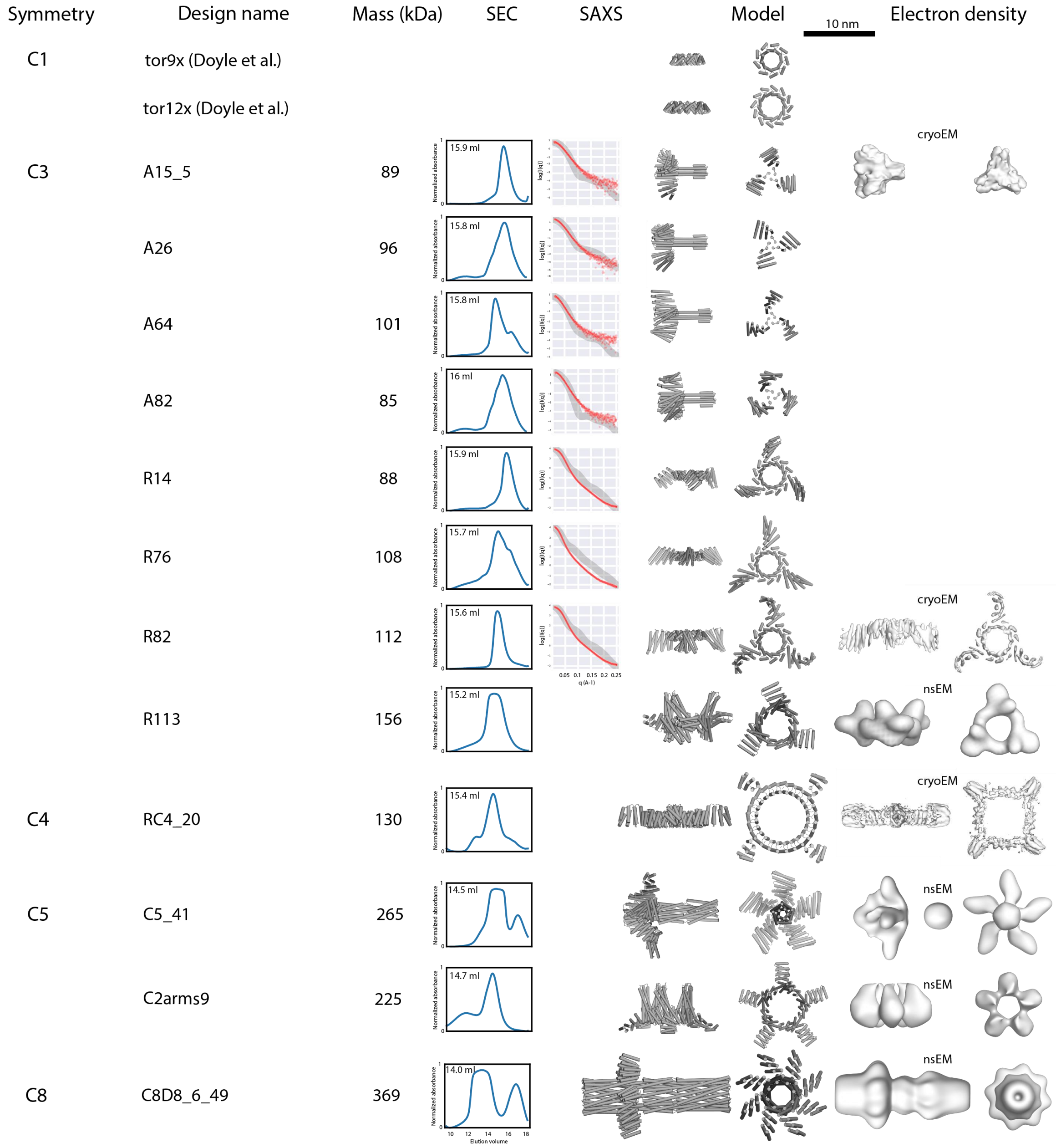


**Figure S2: Detail of the library of axle and ring parts for the design of rotary machines with corresponding symmetry, design nomenclature, oligomeric mass, SEC chromatograms, SAXS traces, designed PDB model, and 3D electron density reconstruction from electron microscopy analysis.** For each SEC trace, the theoretical elution volume corresponding to the correct oligomer state is given in milliliters in black next to the chromatogram. Experimental SAXS traces are shown in the red line, while the theoretical trace corresponding to the design is shown in grey. nsEM: data obtained using negative stain electron microscopy; cryoEM: data obtained using single particle cryoEM.

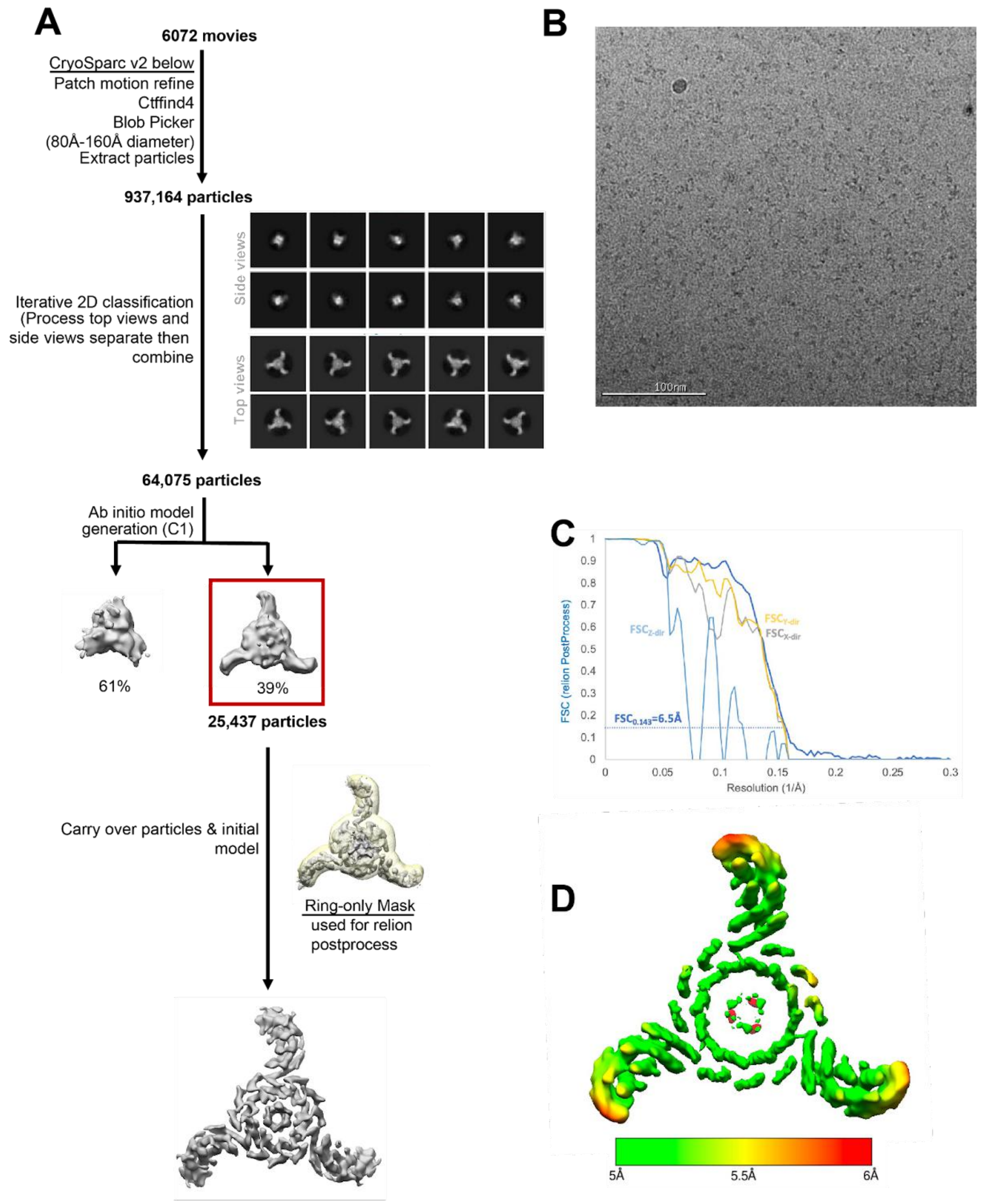


**Figure S3: CryoEM data processing pipelines used to generate electron density and structures of the C3-C3 rotary machine.** (**A**) Detail of the data processing pipeline (**B**) Representative cryoEM micrograph (**C**) FSC validation curve (**D**) Electron density map with corresponding estimated local resolution.

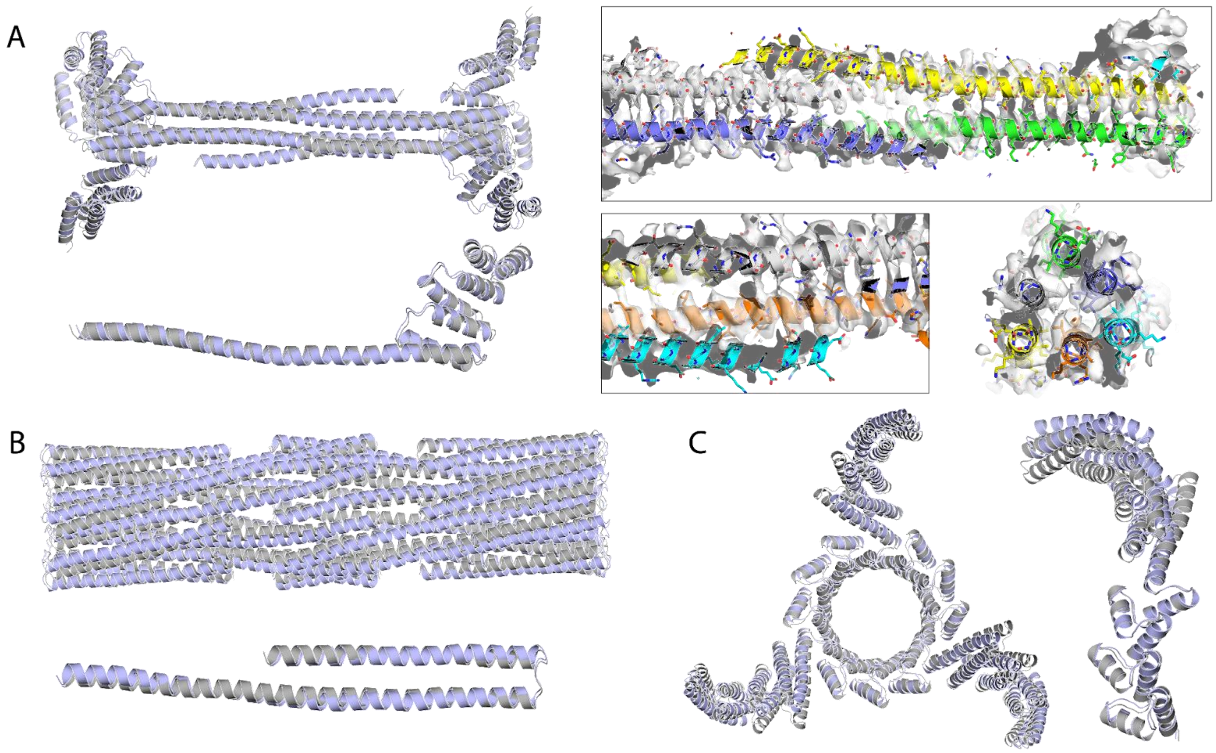


**Figure S4: Detailed comparison of designs versus high resolution cryoEM structures.** The designs were relaxed into experimental cryoEM electron densities using Rosetta FastRelax and SetupForDensityScoring. (**A**) D3 axle design (1552_1na0C3_int2_11); (Left) Superposition of the designed backbone (grey) and backbone relaxed into the experimental electron density (light blue), full structure and single chain alignment. The computed backbone atom RMSD from the designed and experimental structure is 1.930Å. (Right) Detail of side chain density that becomes visible at this resolution (~4Å). (**B**) D8 axle design (D8A_1615). Superposition of the designed backbone (grey) and backbone relaxed into the experimental electron density (light blue), full structure and single chain alignment. The computed backbone atom RMSD from the designed and experimental structure is 2.879Å. (**C**) C3 ring design (R82). Superposition of the designed backbone (grey) and backbone relaxed into the experimental electron density (light blue), full structure and single chain alignment. The computed backbone atom RMSD from the designed and experimental structure is 3.451Å.

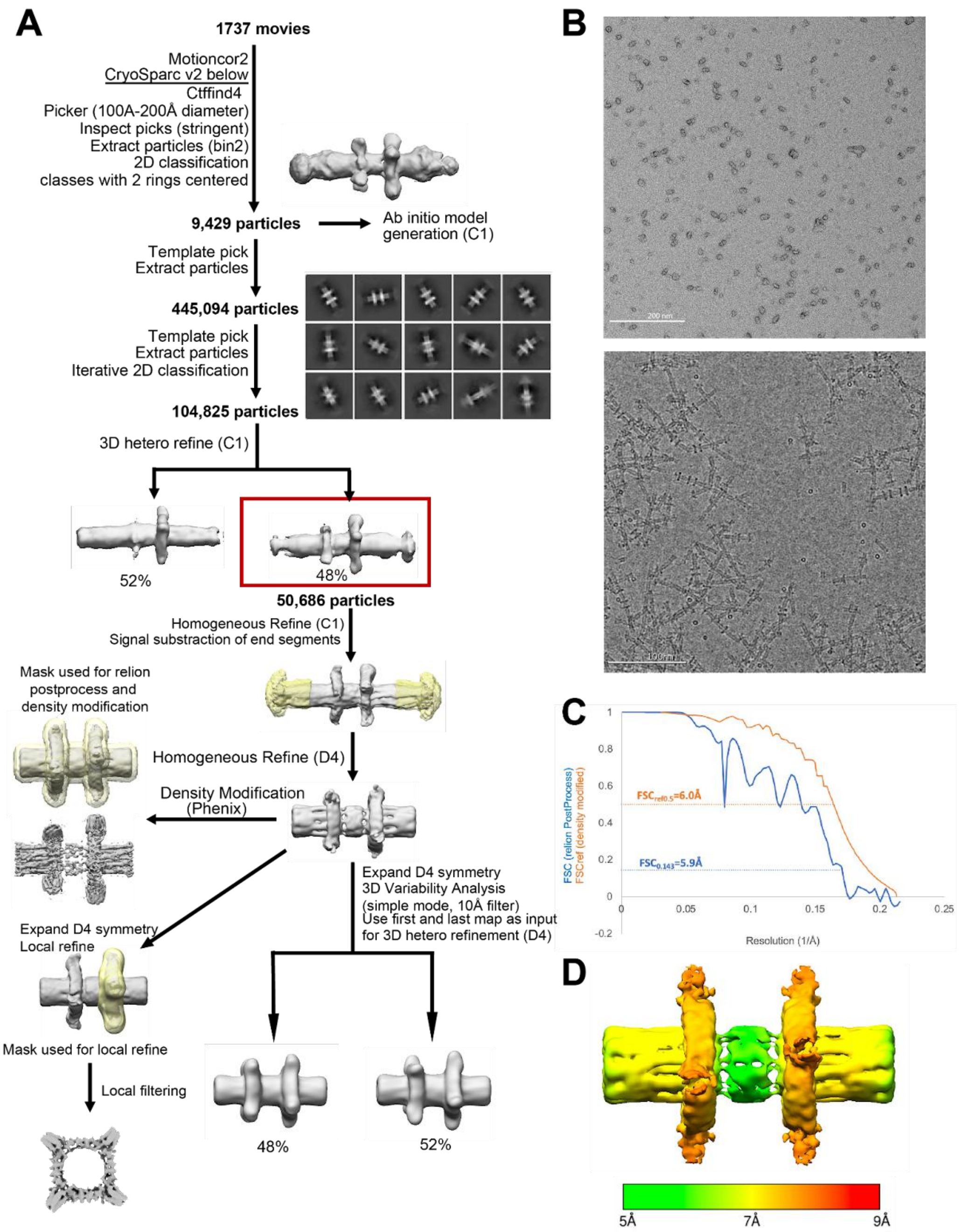


**Figure S5: CryoEM data processing pipelines used to generate electron density and structures of the D8-C4 rotary machine.** Interestingly, this design self-assembled into higher-order fiber-like structures upon freezing, which highlights the unintended effects of cryogenic conditions on protein assemblies. We could however verify that this fiber assembly did not happen at room temperature in solution, as can be seen from SAXS, SEC in figS3, as well as negative stain EM. (**A**) Detail of the data processing pipeline (**B**) Representative cryoEM micrograph. Top: Negative stain; bottom: cryoEM. Freezing conditions seemed to induce fiber formation via end-to-end contact of the D8 axle. (**C**) FSC validation curve (**D**) Electron density map with corresponding estimated local resolution.

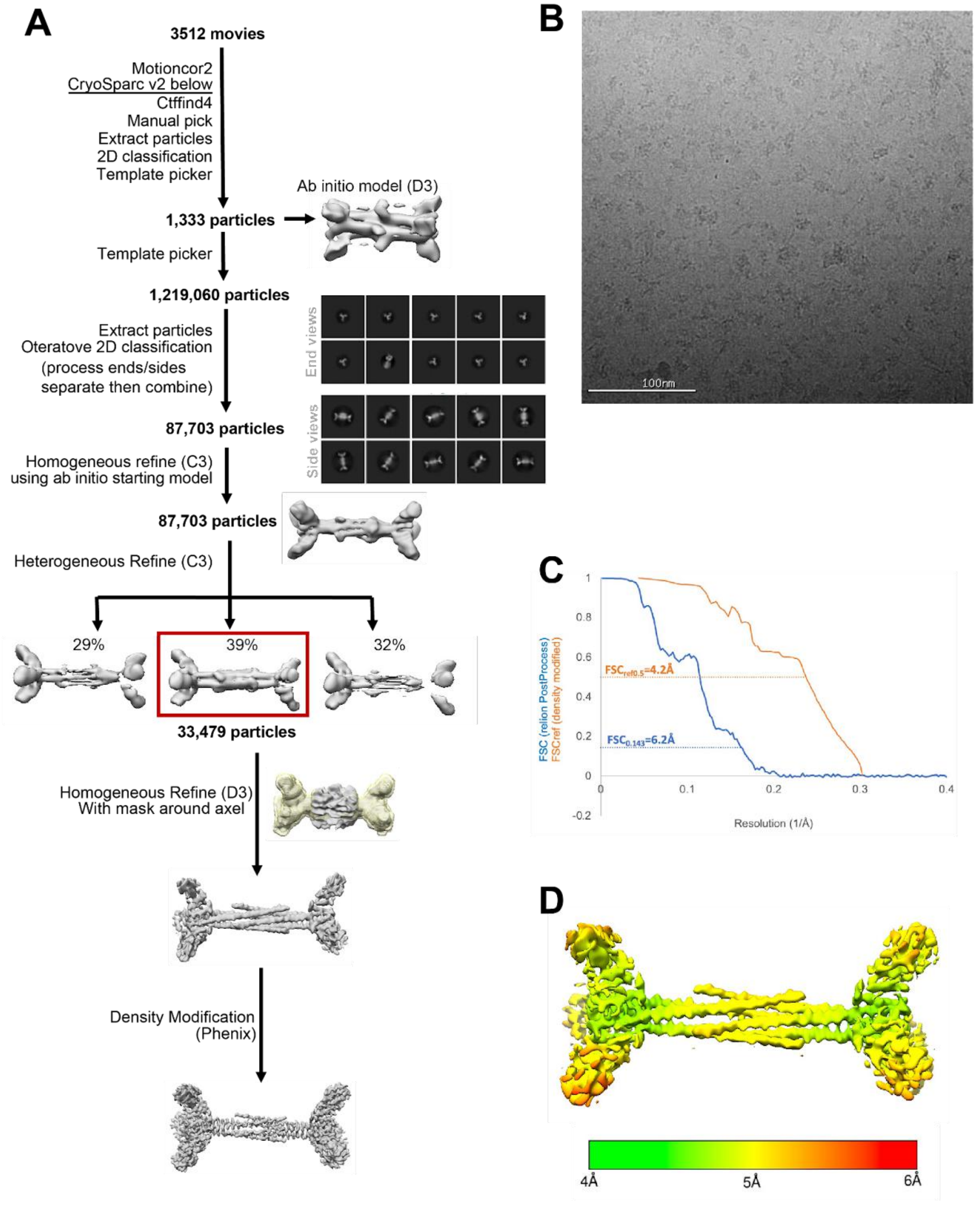


**Figure S6: CryoEM data processing pipelines used to generate electron density and a structure of the D3 axle.** This data was collected on a version of the D3-C5 rotary machine, for which the C5 ring did not have arms extension, thus precluding obtention of clear ring density. We used this dataset to thus focus on obtaining a clear picture of the axle, as detailed here. (**A**) Detail of the data processing pipeline (**B**) Representative cryoEM micrograph (**C**) FSC validation curve (**D**) Electron density map with corresponding estimated local resolution.

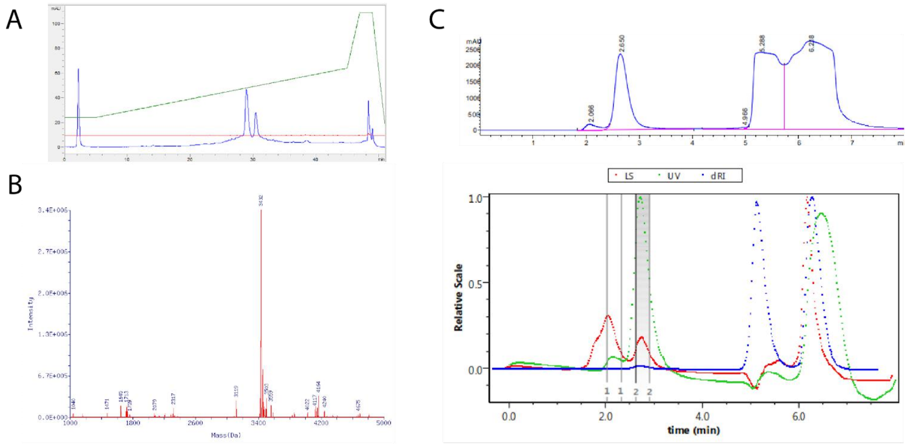


**Figure S7: Chemical synthesis of a 36 residues helical peptide self-assembling into a D3 homohexamer.** (**A**) HPLC chromatogram post synthesis, showing two elution peaks. (**B**) Deconvoluted native mass spectra corresponding to the HPLC peaks. (**C**) Size exclusion chromatography (top, 215nm absorbance) coupled with multiple angle laser light scattering analysis (bottom) of the collected fractions post synthesis and purification. Integration of peak two gives a molecular weight of 25 kDa +/- 7, corresponding to the size of the homohexamer assembly.

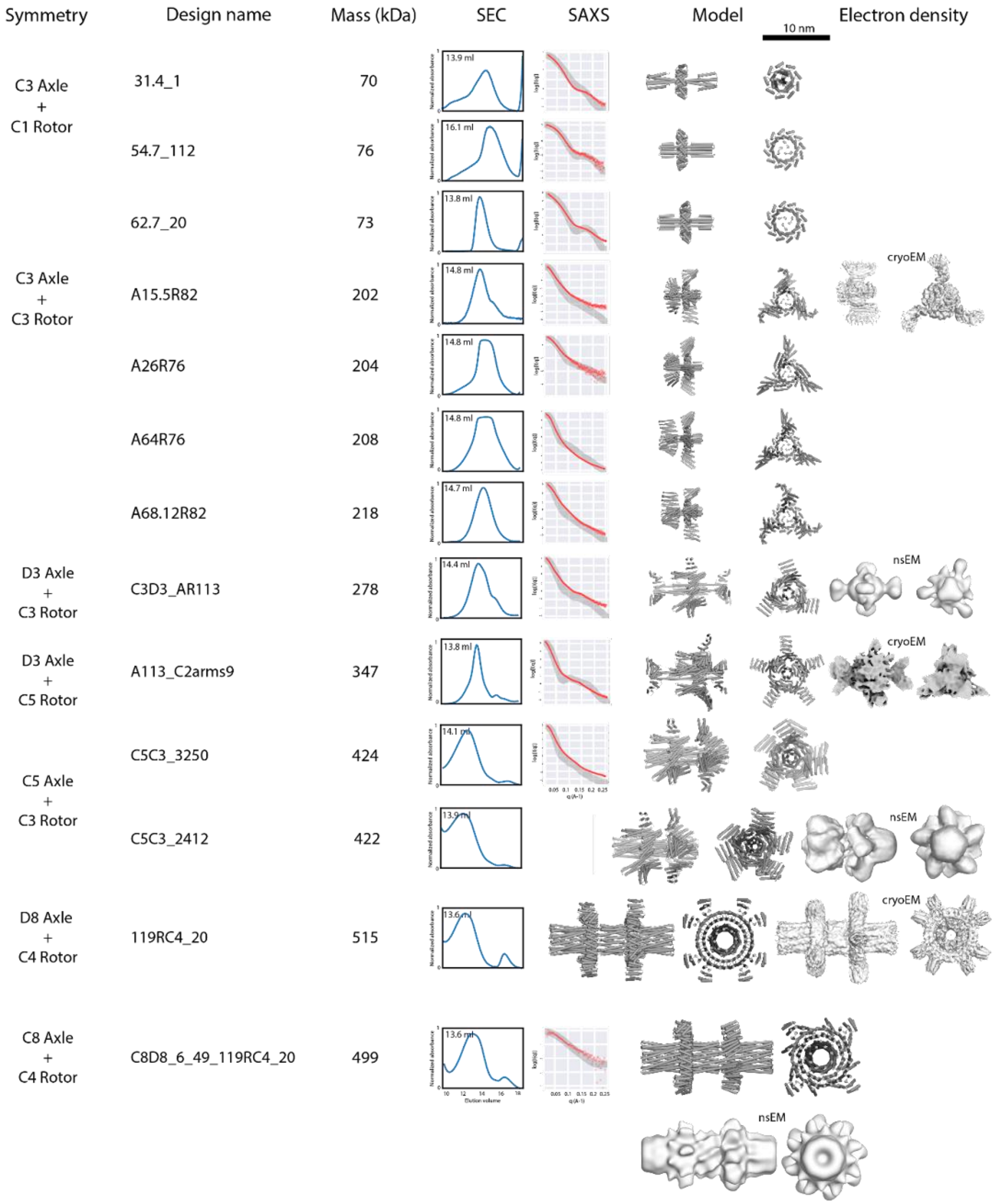


**Figure S8: Detail of the library of fully assembled rotary machines with corresponding symmetry, design nomenclature, oligomeric mass, SEC chromatograms, SAXS traces, designed PDB model, and 3D electron density reconstruction from electron microscopy analysis.** For each SEC trace, the theoretical elution volume corresponding to the correct oligomer state is given in milliliters in black next to the chromatogram. Experimental SAXS traces are shown in the red line, while the theoretical trace corresponding to the design is shown in grey. nsEM: data obtained using negative stain electron microscopy; cryoEM: data obtained using single particle cryoelectron microscopy.

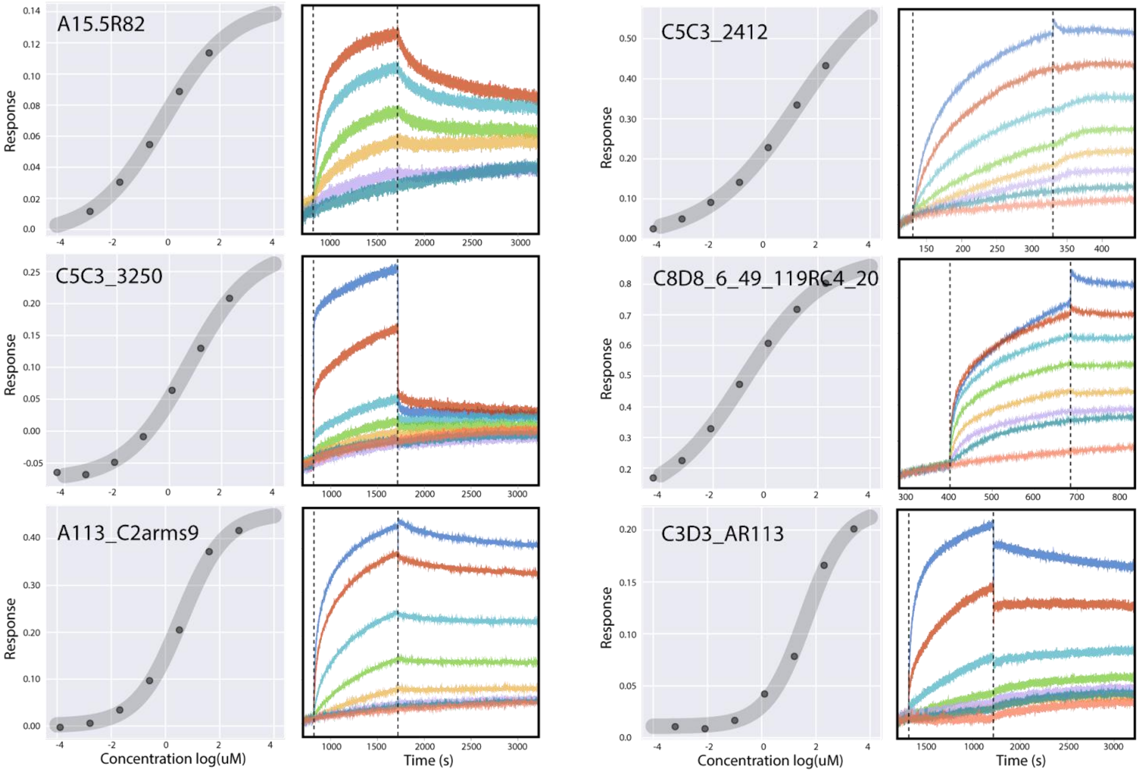


**Figure S9: Biolayer interferometry assays measuring *in vitro* assembly of ring and axle parts into a full rotary system.** For each 6 designs, the equilibrium binding curves from biolayer interferometry binding assays is shown on the left and the corresponding Biolayer interferometry kinetic binding traces shown on the right. Biotinylated axles were immobilized on the tip and the binding in a solution of fre rings was measured. For both D3-C5 and D3-C3, a fresh ring solution was prepared by buffer exchange from citrate buffer to TBS with reducing agent, and immediately used for binding assays.

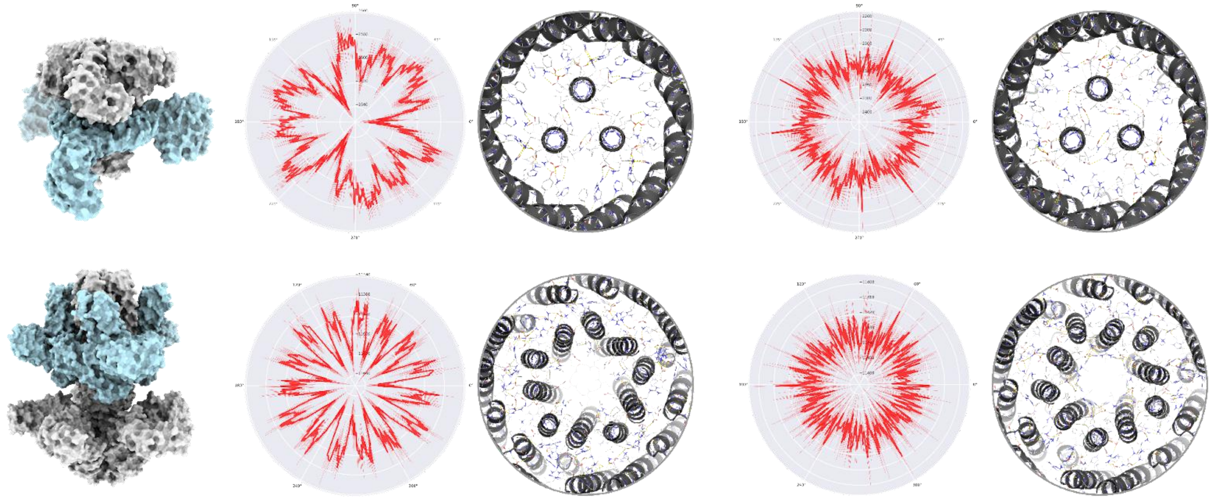

**Figure S10: Example of designed energy landscapes. The shape, periodicity and energetics differ drastically depending on the residue identities and contact types for the same protein scaffold.** (Top) Two C3-C3 rotors design trajectories; (Bottom) Two C5-C3 rotors design trajectories: C5C3_3250, C5C3_2412; (Left) PDB models (ring color in cyan and axle in grey); (Right) Energy landscapes shown as polar maps, depicting Rosetta Energy Units (REU) vs rotation angle generated by sampling along the rotational degree of freedom while using Rosetta relax, minimization of side chains and scoring for each rotation bins. The mean energy landscape obtained from 10 independent trajectories is shown in red with error bars depicting the standard deviation. The designed interface between axle and ring at angle=0 is shown beside as cross-sections, showing residue identities and contacts and hydrogen-bond networks (yellow dashed lines) with the helical backbone (shown in grey).

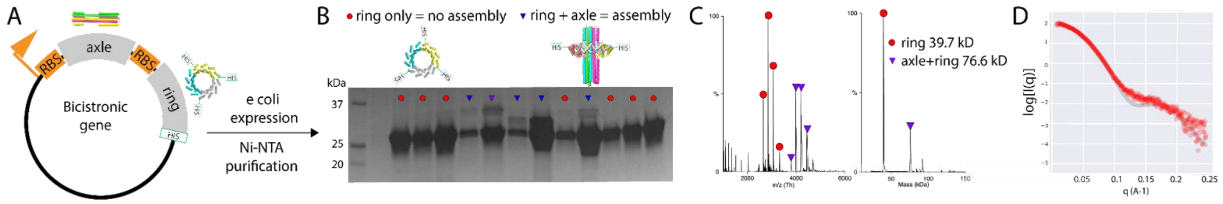


**Figure S11:** ***In vivo* assembly of two component rotary machines from bicistronically expressed axle and ring parts** (**A**) Plasmid architecture for the bicistronic expression system based pET29b+. (**B**) SDS-PAGE after Ni-NTA purification while bicistronically expressing axle and ring in the same cell. A single band indicates that the ring did not pull down the axle (lanes marked with red circle), while 2 bands indicate assembly of axle and ring (marked as an triangle, purple triangle marks the selected design) (**C**) Convoluted and deconvoluted native mass spectrums of the isolated C3-C3 rotary machine. (**D**) SAXS traces of the purified protein. The experimental trace is shown in the red line, while the theoretical trace corresponding to the design is shown in grey.

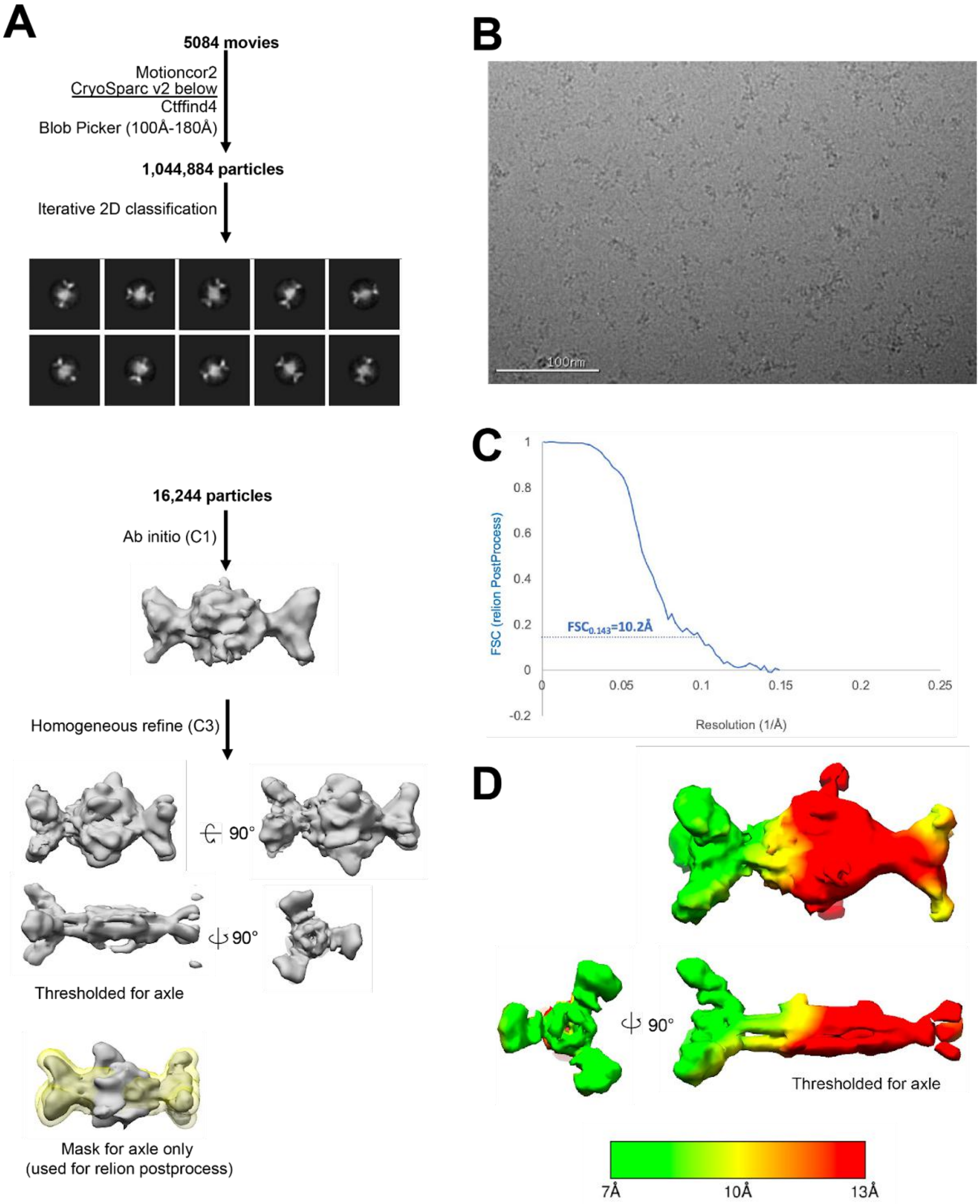


**Figure S12: CryoEM data processing pipelines used to generate electron density and structures of the D3-C3 rotary machine.** (**A**) Detail of the data processing pipeline. The C1 reconstruction in stain yielded C3 features, which allowed us to further process the design with C3 symmetry imposed here. The C3 reconstruction also showed ring density that was polar, consistent with the rotor design. The ring density looks very similar when processed in D3, while yielding a better model for the axle (which has D3 symmetry by design). Therefore we used a whole model processed in D3 mode to present in figure 3C, which is closest to the actual symmetry and structure of the model (**B**) Representative cryoEM micrograph (**C**) FSC validation curve for C3 reconstruction. (**D**) Electron density map with corresponding estimated local resolution.

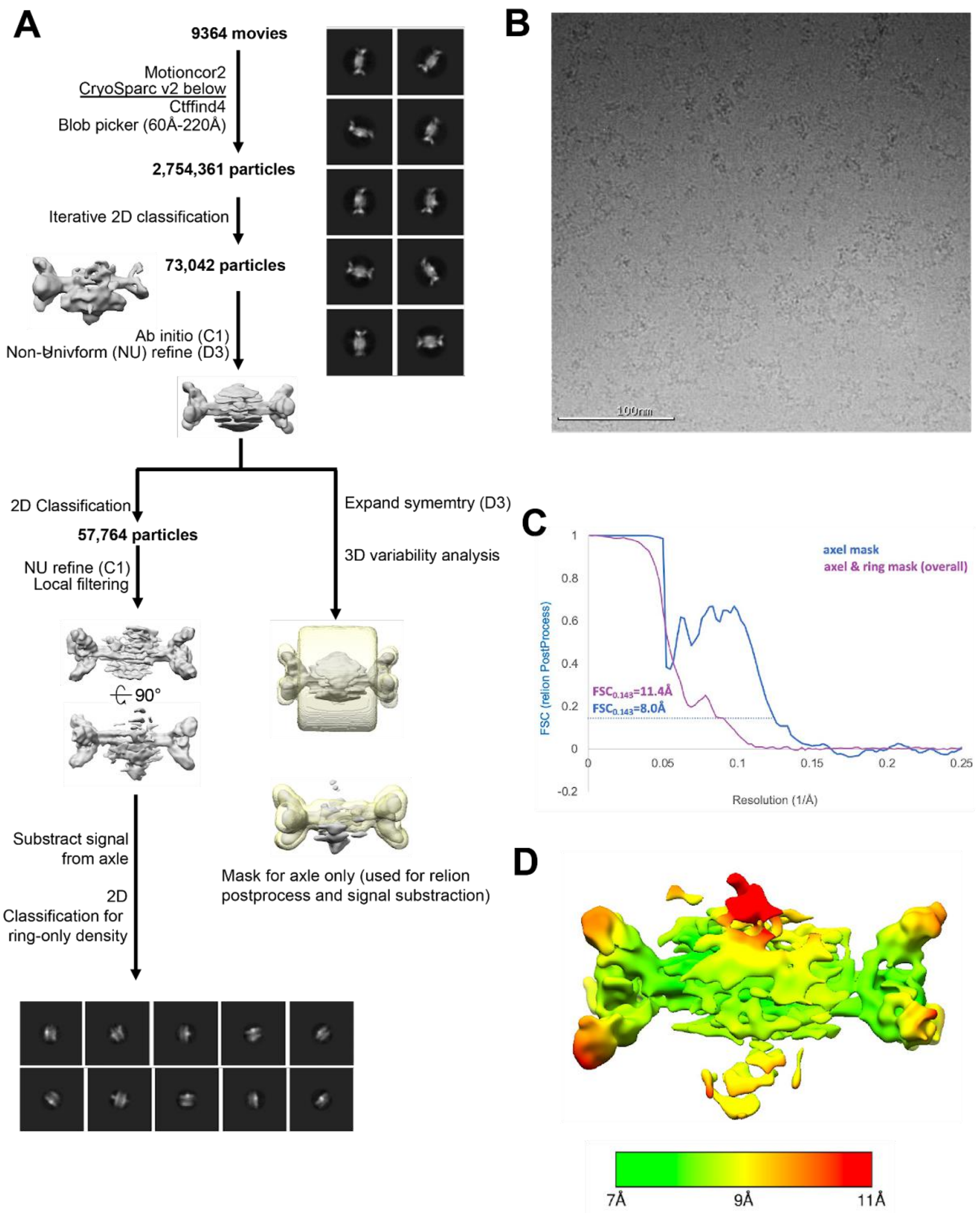


**Figure S13: CryoEM data processing pipelines used to generate electron density and structures of the D3-C5 rotary machine.** (**A**) Detail of the data processing pipeline (**B**) Representative cryoEM micrograph (**C**) FSC validation curve of C1 reconstruction (**D**) Electron density map with corresponding estimated local resolution.

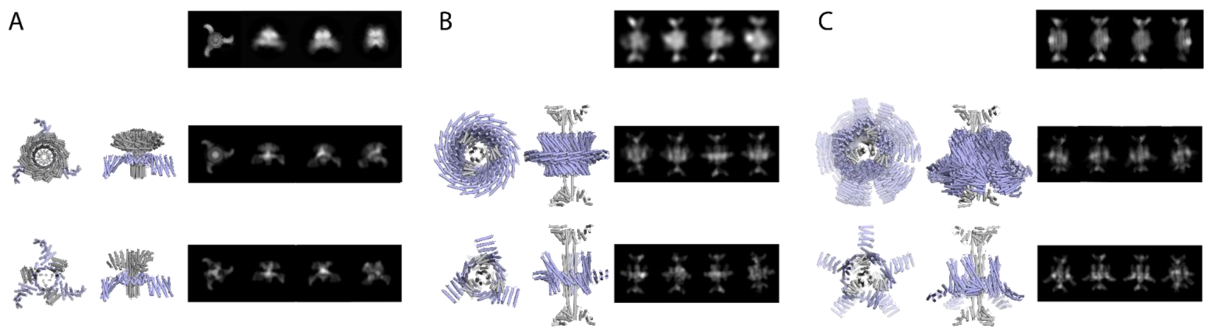


**Figure S14: Detail of DOF sampling for the generation of the theoretical cryoEM 2D class averages projections compared to experimental data and models.** (**A**) C3-C3 rotor (top) Experimental 2D averages (middle) Projections obtained when taking into account the rotational DOF and simulating 10 trajectories with corresponding PDB model shown on the left (bottom) Projections obtained when not taking into account the rotational DOF with corresponding PDB model shown on the left (**B**) D3-C3 (top) Experimental 2D averages (middle) Projections obtained when taking into account the rotational DOF and simulating 10 trajectories with corresponding PDB model shown on the left (bottom) Projections obtained when not taking into account the rotational DOF and simulating 10 trajectories with corresponding PDB model shown on the left (**C**) D3-C5 rotor (top) Experimental 2D averages (middle) Projections obtained when taking into account the rotational and translational DOF and simulating 10 trajectories with corresponding PDB model shown on the left (bottom) Projections obtained when not taking into account the rotational and translational DOFs and simulating 10 trajectories with corresponding PDB model shown on the left.

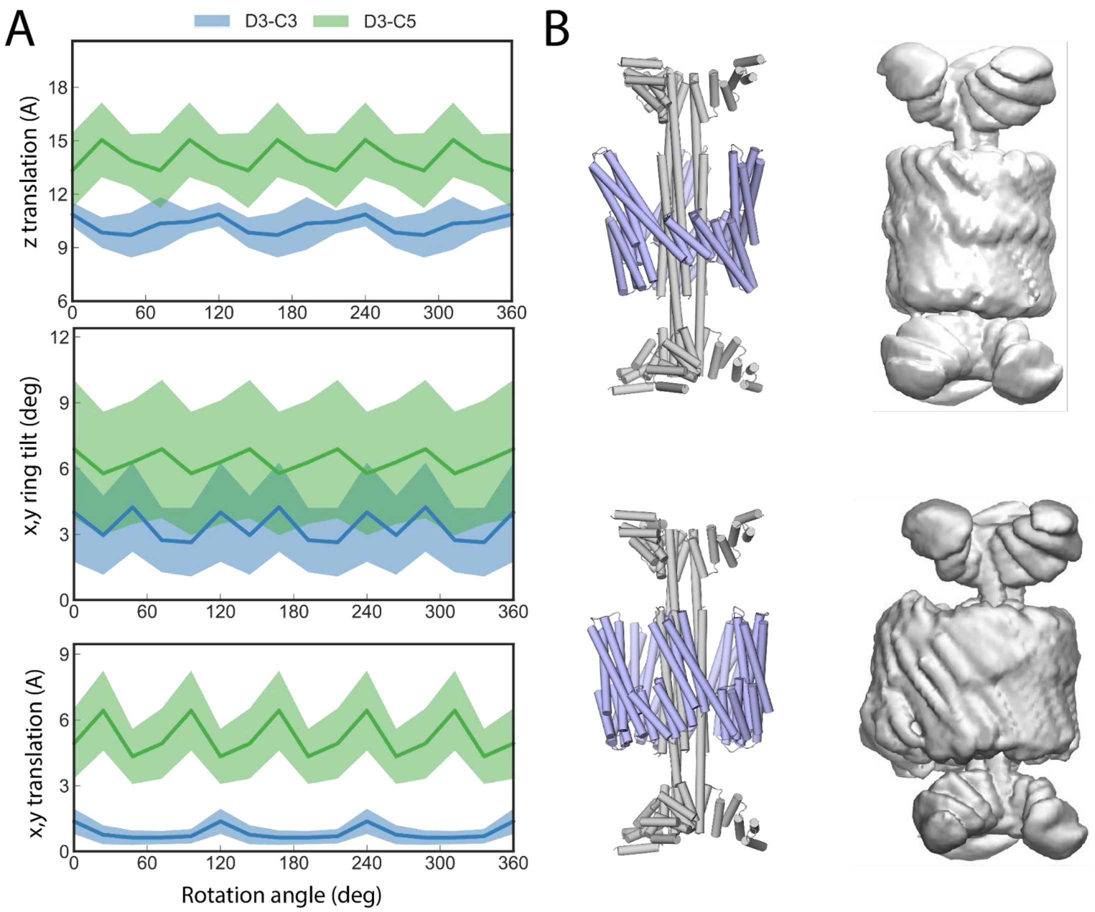


**Figure S15: Molecular dynamics simulations performed on D3-C3 and D3-C5 rotary machine assemblies to investigate the DOF of motion** (**A**) The interface shape, size and symmetry of these two design results in different DOFs: the D3-C3 was found to rotate along the z axis, while the D3-C5 ring showed rotation along x, y and z, as well as translation in z and y. The top panel shows z translation of rings relative to rotation, the middle panel shows the x and y rotation or the ring, or tilt, relative to rotation and the bottom panel shows the x, y translation of the ring relative to the rotation around the axle. (**B**) Top: D3-C3; Bottom: D3-C5; Left: PDB models; Right: density maps of the backbone atoms showing averaged motion of ring and axle relative to each other.

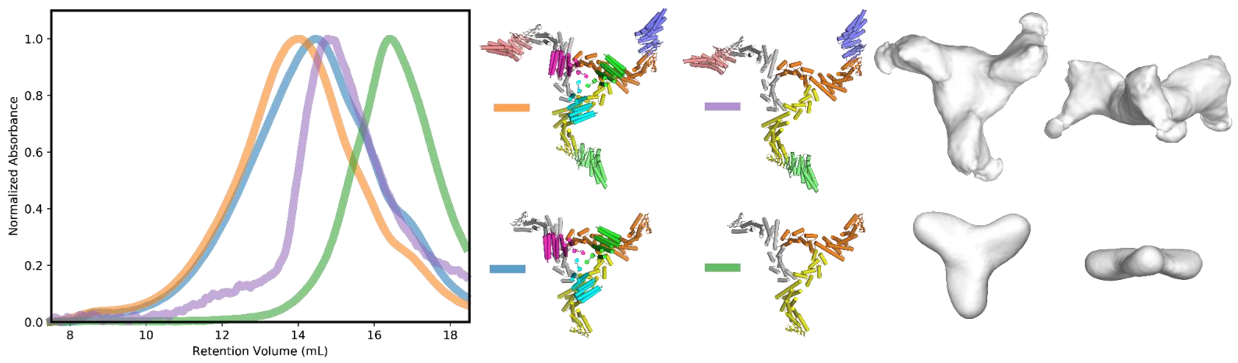


**Figure S16: Rotary machine modular extension by systematic fusion with reversible heterodimers.** (Left) SEC elution profiles corresponding the C3-C3 rotor assembly with and without heterodimer arms extension, or ring only with or without heterodimer extension. (Right) Top and side views of negative stain 3D reconstruction corresponding to the ring only with or without the heterodimer arms.

| | C3-C3 | D3 axle | D3-C5 | D3-C3 | D8-C4 |
|---|---|---|---|---|---|
| **Number of micrographs** | 6,072 | 3,512 | 9,364 | 5,084 | 1,737 |
| **Nominal magnification** | 130,000X | | | | 36,000X |
| **Voltage** | 300 kV | | | | 200 kV |
| **Electron Fluence** | 90e-/Å$^2$ | | | | 65e-/Å$^2$ |
| **Pixel size** | 1.05Å | | | | 1.16Å |
| **Defocus range** | -1.7 to -2.8µm | -1.5 to -2.4µm | -1.3 to -2.5µm | -0.5 to -3.8µm | -1.4 to -4.2µm |
| **EMDB ID** | | | | | |
| **Map resolution 0.143 FSC** | 6.5Å | 6.2Å | 8Å | 10.2Å | 7.2Å |
| **Density Modified Resolution 0.5Ref** | n/a | 4.2Å | n/a | n/a | 7.0Å |
| **Symmetry Imposed** | C3 | D3 | C1 & D3 | C3 | D4 |
| **Number of particles** | 25,437 | 33,479 | 73,042 (D3)<br>57,764 (C1) | 16,244 | 50,686 |
| **Refinement** | | | | | |
| **Map Sharpening B Factor** | -330Å$^2$ | -284Å$^2$ | -471Å$^2$ | -503Å$^2$ | -430Å$^2$ |

**Table S1. Cryo-EM data collection and Refinement statistics**

**Movie S1: 3D Variability analysis (3DVA) first component of the D3-C5 design.** 3DVA was done in cryosparc v2 after D3 symmetry expansion while using default processing parameters and a 5Å low-pass filter. Variability results are displayed as the first trajectory component of variability calculated in simple mode as a 20 frame output, which are low-pass filtered to 5Å for clarity. Volume series is rendered as a movie using UCSF chimera. Movie shows rotation of the rotor through the volume series, as well as a transverse cross section of the central ring and axle.

**Movie S2: 3D Variability analysis (3DVA) second component of the D3-C5 design.** 3DVA was done in cryosparc v2 after D3 symmetry expansion while using default processing parameters and a 5Å low-pass filter. Variability results are displayed as the second trajectory component of variability calculated in simple mode as a 20 frame output, which are low-pass filtered to 5Å for clarity. Volume series is rendered as a movie using UCSF chimera. Movie shows rotation of the rotor through the volume series, as well as a transverse cross section of the central ring and axle.

**Movie S3: 3D Variability analysis (3DVA) third component of the D3-C5 design.** 3DVA was done in cryosparc v2 after D3 symmetry expansion while using default processing parameters and a 5Å low-pass filter. Variability results are displayed as the third trajectory component of variability calculated in simple mode as a 20 frame output, which are low-pass filtered to 5Å for clarity. Volume series is rendered as a movie using UCSF chimera. Movie shows rotation of the rotor through the volume series, as well as a transverse cross section of the central ring and axle.

**Movie S4: 3D Variability analysis (3DVA) of the D3-C3 design.** 3DVA was done in cryosparc v2 after D3 symmetry expansion while using default processing parameters and a 5Å low-pass filter. Variability results are displayed as the first trajectory component of variability calculated in simple mode as a 20 frame output, which are low-pass filtered to 5Å for clarity. Volume series is rendered as a movie using UCSF chimera. Movie shows rotation of the rotor through the volume series, as well as a transverse cross section of the central ring and axle.

**Movie S5: 3D Variability analysis (3DVA) of the D8-C4 design**. 3DVA was done in cryosparc v2 after D4 symmetry expansion while using default processing parameters and a 10Å low-pass filter. Variability results are displayed as the second trajectory component of variability calculated in simple mode as a 10 frame output, which are low-pass filtered to 15Å for clarity. Volume series is rendered as a movie using UCSF chimera. First and last frames of the second trajectory component were used as input for downstream refinement of distinct structures.
